# Multivalent weak contacts shape chaperone-nascent protein interactions

**DOI:** 10.64898/2026.03.09.709851

**Authors:** Nandakumar Rajasekaran, Dmitri Toptygin, Ting-Wei Liao, Vincent J. Hilser, Taekjip Ha, Christian M. Kaiser

## Abstract

Molecular chaperones interact with non-native proteins, playing crucial roles in preventing misfolding and enable efficient folding in the cellular environment. Trigger factor is a bacterial chaperone that binds to ribosomes, interacting with nascent polypeptides emerging from the ribosome and guiding their early folding steps. In contrast to the central role of the chaperone in promoting folding of newly synthesized proteins, its dynamic interactions with nascent chains emerging from the ribosome remain poorly understood. Here, we use single-molecule fluorescence and optical tweezers approaches to directly observe and characterize trigger factor interactions with a ribosome-bound client protein at increasing chain lengths. We find that trigger factor binding to nascent proteins is best described by a combination of multiple weak, dynamic interactions that are established after the chaperone docks onto the ribosome and evolve during polypeptide elongation. Application of mechanical force perturbs trigger factor binding, supporting a multivalent interaction model. This binding mode may help to stabilize nascent proteins against misfolding while allowing them to dynamically sample conformational space in search of their native structures.

## Introduction

The biological functions of cellular proteins are sustained by their native structures. While a small fraction of the proteome can spontaneously fold into native structures (1), the vast majority of extant proteins require the assistance of molecular chaperones to properly fold and avoid aggregation (2–5). Early folding begins as nascent proteins emerge from the ribosome during synthesis, which is particularly important for large, multi-domain proteins (6). The bacterial chaperone trigger factor associates with ribosomes (7) and, together with the Hsp70 chaperone DnaK, assists in the early stages of folding (8, 9).

Originally identified as a factor that maintains outer membrane proteins translocation-competent (10), trigger factor was subsequently recognized as a general folding catalyst (11). Trigger factor binds relatively weakly to both non-translating ribosomes and isolated proteins but has increased affinity for ribosome-associated nascent polypeptides (12, 13). Consistent with acting at early stages during protein synthesis, it binds to nascent proteins that are not yet structured, delaying their folding (9) and reducing misfolding (14). Recent studies indicate that the chaperone can also interact with non-native regions in partially folded structures (15) and may even accelerate folding (16) of nascent proteins.

Global analysis by ribosome profiling indicated that trigger factor stably binds to nascent chains of 100 or more amino acids in length (17). Length- and sequence-dependent binding is further supported by biochemical experiments measuring overall affinity of the chaperone for ribosome-bound nascent chains of cytosolic proteins (18, 19). Structural studies of isolated client protein fragments using NMR spectroscopy have identified four binding sites, distributed across several domains of trigger factor, that can engage with individual interaction sites in the client protein (20). However, the detailed kinetics of this multivalent interaction in the context of translating ribosomes have not yet been defined.

To shed light on the precise nature of the interactions between trigger factor and nascent chains, we developed single-molecule assays to dissect binding of the chaperone to an authentic nascent client protein. We observed transient binding of trigger factor to nascent proteins with dwell times that varied non-monotonically with nascent chain length. The kinetics of trigger factor interactions reveal a complex multi-phasic mechanism defined by the properties of the nascent protein. Perturbations of trigger factor-nascent chain complexes with mechanical force weakens these interactions, confirming a multi-valent binding mode in the context of the ribosome. Combined with modeling the properties of nascent proteins, our experiments resolve the interactions that ultimately underpin chaperone function.

## Results

### Single-molecule TIRF microscopy resolves transient trigger factor-RNC interactions

To directly monitor the dynamic interaction of trigger factor with ribosome-nascent chain complexes (RNCs), we developed a single-molecule dual-color TIRF microscopy approach (Fig. 1A) employing elongation factor G (EF-G), which had previously been demonstrated to be an authentic client protein of the chaperone (14, 21). The N-terminal GTPase-domain (G-domain) of EF-G encompasses 293 amino acids and remains unstructured until it is fully extruded from the ribosome during synthesis (22), providing an opportunity to characterize the interaction of a genuine trigger factor client nascent protein at multiple chain lengths without the complication of folding.

**Figure 1.**
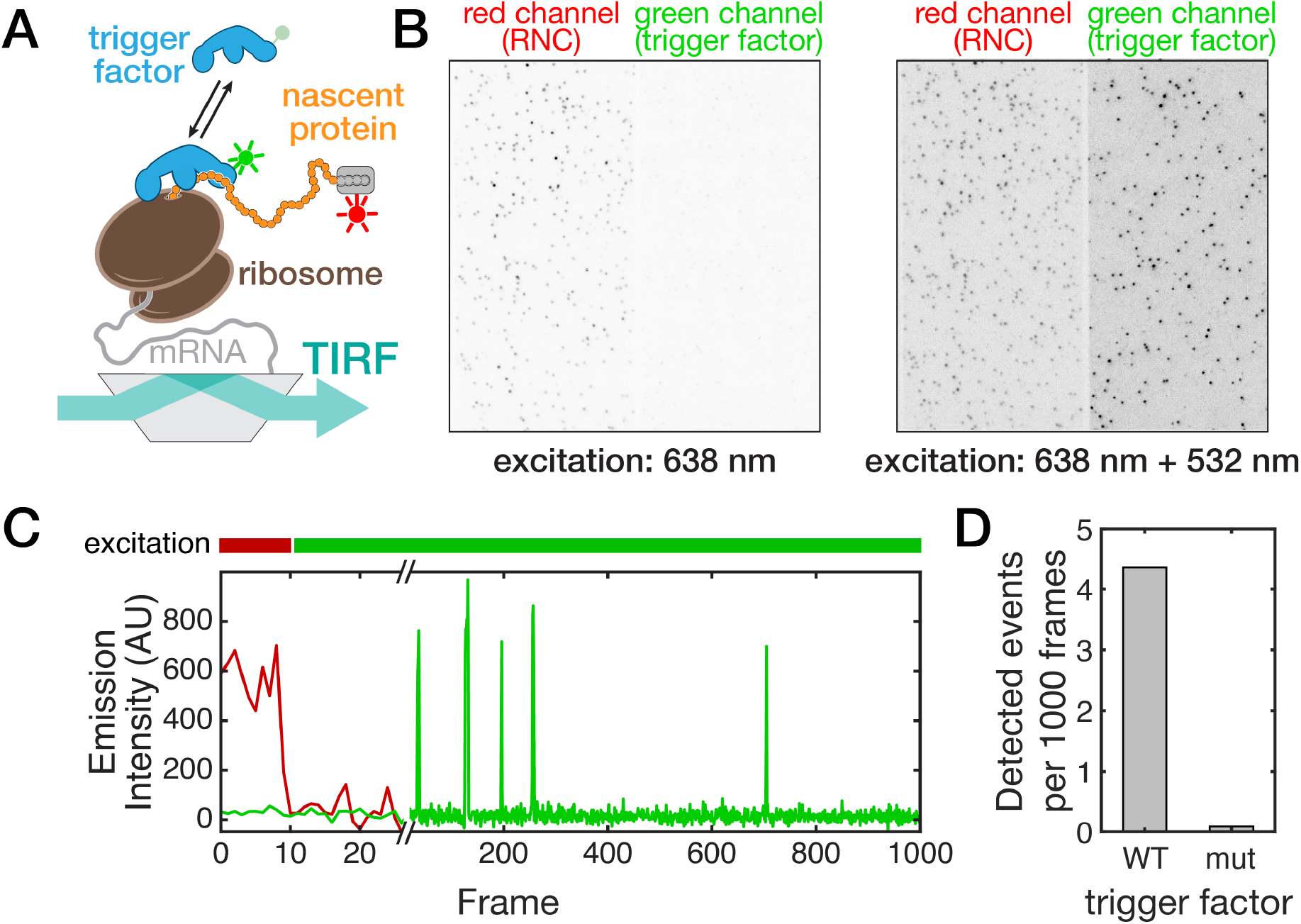
Single-molecule observation of chaperone-nascent chain interactions. **A**. Schematic of the experimental setup. Ribosome-nascent protein complexes (RNCs) are surface-immobilized via the messenger RNA (mRNA). TIRF illumination allows detection of Atto532-labeled trigger factor molecules upon RNC binding. The nascent protein is labeled with sulfo-Cy5-derivatized SpyCatcher (grey) at an N-terminal SpyTag. **B**. Time-integrated wide-field TIRF images showing the initial excitation with 638-nm laser (left, 10 frames) and the entire movie after excitation with 532-nm laser (right, 30 frames). Initial excitation is used to identify the positions of individual immobilized RNCs (red channel on the left). Dots in the green channel (right) that co-localize with the red dots represent RNC-bound TF molecules. **C**. Example trajectory of fluorescence emission intensity versus time at the position of an RNC, which is detected by red illumination during the initial 10 frames. Subsequently, red illumination is turned off, and green illumination is turned on. Individual TF binding and dissociation events are detected as spikes in the recorded fluorescence emission intensity. Frame rate: 10 Hz. **D**. Comparison of binding events detected with wild-type trigger factor (WT) and a mutant defective in ribosome binding (mut), demonstrating specific binding to surface-immobilized RNCs.

To resolve chaperone-nascent chain interactions at defined nascent chain lengths, representing snapshots of protein synthesis, we generated RNCs by *in vitro* translation of non-stop messenger RNAs (mRNAs) (14). RNCs are stably stalled at the 3’-terminus of these non-stop mRNAs. We first produced a nascent chain encompassing the first 135 amino acids (aa) of the EF-G sequence, termed here RNC135. Selective ribosome profiling studies indicated that trigger factor strongly engages nascent chains with lengths above 100 amino acids (17, 23). We thus expected that the chaperone binds robustly to RNC135.

Isolated RNC135 molecules were immobilized on streptavidin-coated glass slides by means of a biotinylated oligonucleotide hybridized to the mRNA near the 5’-terminus (24). An N-terminal SpyTag (25) served to label the nascent chain with sulfo-Cy5-derivatized SpyCatcher (Fig. 1A) to visualize surface-immobilized ribosome-nascent chain complexes, which appeared as individual spots in the red channel (Fig. 1B, left panel). Subsequent excitation of Atto532-labeled trigger factor resulted in the transient appearance of spots in the green channel that mostly co-localized with the detected RNC signals (Fig. 1B, left panel). Typically, around 200 fluorescent spots were detected in the field of view. The Cy5 signal was detected only in the red detection channel (Fig. 1B), confirming spectral separation. Control experiments (Suppl. Fig. S1) confirmed that the observed fluorescent spots were due to specific surface-immobilization of RNCs, ruling out artifacts such as non-specific immobilization of RNCs or released nascent chains that might interact with trigger factor.

To resolve interaction dynamics, we monitored binding events at individual spots at a frame rate of 10 Hz. At the beginning of each measurement, the sample was illuminated with the 638-nm laser for 1 s to detect the positions of RNCs. The 638-nm laser was then turned off, and the sample was illuminated with the 532-nm laser to image trigger factor. Fluorescence intensities at each identified RNC position were transformed into time series of green and red intensities (Fig. 1C). The time traces showed brief recurring spikes in green fluorescence, reflecting dynamic interactions of trigger factor with RNC135.

Co-localization of the sulfo-Cy5 (RNC135) and Atto532 (trigger factor) signals indicate specific association of the chaperone with surface-immobilized RNCs. In control experiments with a trigger factor mutant deficient in ribosome-binding, in which three residues (FRK) in the ribosome-binding domain are replaced with alanine (AAA) (7), we observed a reduction in binding events by 98% (4.36 and 0.09 events at each RNC per 1000 frames, respectively; Fig. 1D). These results rule out possible artifacts arising from non-specific sticking of the chaperone to the glass surface and indicate that ribosome-binding is necessary for detecting trigger factor interaction in our experiments.

### Nascent chain interactions stabilize trigger factor-RNC interactions

To quantitatively characterize the kinetics of trigger factor-RNC interactions, we determined the duration (dwell times) of individual binding events (Fig. 2A, see Methods for details). The survival function *S*(*t*) provides a good visual representation of the dwell time variability (Suppl. Fig. S2). For trigger factor bound to RNC135, *S*(*t*) falls from 1 to 1/*e* at *t*_*1/e*_ = 1.356 s. For an exponential survival function, t_1/e_ would represent the mean dwell time and correspond to an off rate of 0.74 s^−1^. By analyzing the time between binding events and assuming second-order binding kinetics, we estimate an association-rate constant of 26.3 µM^−1^ s^−1^ (Suppl. Fig. S3). *In vitro* ensemble measurements of 75 amino acid-long nascent chains from two unrelated proteins, HemK and proOmpA, yielded dwell times of 1.1 s and 1.7 s, respectively (13), comparable to the value obtained here. The rate constants for association reported for these nascent chains (240 and 200 µM^−1^ s^−1^) are higher than the value we obtained, which may reflect differences in the nature of the client proteins. The kinetic parameters we determine here for RNC135 correspond to a dissociation constant of K_D_ = 28.1 nM, within the range of previously reported K_D_ values for TF-RNC interaction (ranging from 2 nM (13) to 500 nM (26)). Overall, the values determined here are consistent with those obtained by other methodologies, validating our approach.

**Figure 2.**
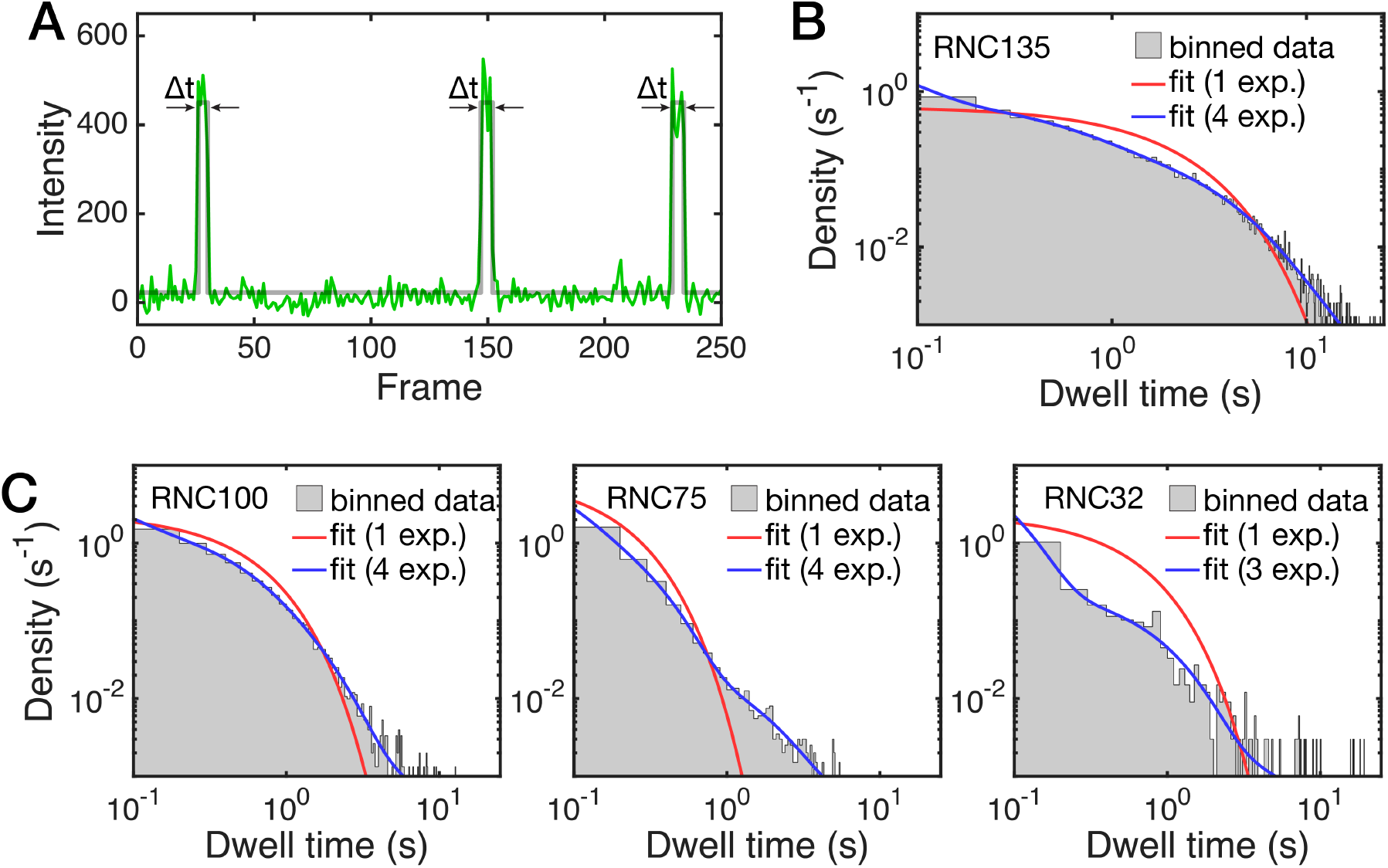
Chaperone-RNC interactions show multi-phasic kinetics. **A**. Fluorescence time trajectory (green) and state assignment (grey line) for RNC135. This segment shows three binding events with their respective dwell times Δt. **B**. Probability density of chaperone dwell times for RNC135. A single-exponential model (red line) does not describe the binned dwell times (grey area) well, whereas adequate fits are obtained with a multi-exponential model (blue line) containing 4 terms. **C**. Binned dwell times and fits for RNCs with decreasing chain lengths, plotted as in panel B.

Trigger factor binds to both the ribosome and the nascent chain. To dissect the contributions of these interactions, we generated RNCs with shorter nascent chains containing 100 and 75 residues of EF-G (RNC100 and RNC75) plus an N-terminal SpyTag for fluorescent labeling (Fig. 1A). For RNC100, the survival function *S*(*t*) reaches 1/*e* at t_1/e_ = 0.43 s and drops to t_1/e_ = 0.17 s for RNC75. Further shortening the nascent chain to 32 EF-G residues yields t_1/e_ = 0.087 s. At this chain length, the EF-G sequence is fully sequestered in the exit tunnel and is thus not be accessible for binding by trigger factor. The N-terminal SpyTag used to label the nascent protein with fluorescent SpyCatcher is extruded from the ribosome in RNC32 complexes. Potentially, the label may interact with the chaperone and affect the measured dwell times. We generated a control construct (RNC_control_) that contains only the N-terminal 25 amino acids corresponding to the SpyTag/linker sequence used for labeling, but no EF-G residues (RNC_control_). The nascent protein in RNC_control_ is fully sequestered inside the exit tunnel. To label RNC_control_, we utilized SpyTagged ribosomes (27). RNC_control_ exhibits a survival function that drops to 1/e at t_1/e_ = 0.084 (Suppl. Fig. S2), very similar to that of RNC32 and consistent with values for non-translating ribosomes determined previously in vitro (13) and in cells (28). The N-terminal label, therefore, does not appear to affect the measured dwell times. Taken together, our measurements are consistent with previous reports indicating that the stability of trigger factor-nascent chain interactions begins to increase significantly at chain lengths of approximately 100 amino acids on average (17, 23).

### Multiple interactions stabilize trigger factor interactions with long nascent chains

The survival function for trigger factor-RNC135 interaction appears to follow an exponential law. However, the dwell time distribution is not well described by a single exponential term (Fig. 2B, red line). Due to the large number of observed events (12976 events in the RNC135 data set), we were able to confidently resolve multiple exponential terms. Based on the Akaike information criterion (see Suppl. Table 1 for details), we determined that four exponential terms are required to obtain an adequate fit for RNC135 (Fig. 2B, blue line). Generally, four-exponential binding kinetics requires a system containing at least five states (one free state and at least four different bound states). Our analyses thus suggest that several interactions stabilizing the chaperone-RNC complex form and break independently, in line with NMR experiments that showed four distinct, low-affinity binding sites in trigger factor (20). Dissociation of trigger factor is observed in the TIRF experiments only when all interactions with the nascent protein and with the ribosome are dissolved simultaneously.

The results described above indicate that a set of transient interactions contribute to the binding of trigger factor to RNCs. To determine how the stability of the chaperone-RNC complex evolves as the nascent chain is extended, we experimentally determined the dwell times of several additional RNCs with nascent chain lengths up to 325 amino acids, corresponding to extrusion of almost the full G-domain of EF-G. All samples produced dwell times that are described by at least four exponential terms (Suppl. Fig. S4).

To quantify the kinetic stability of trigger factor interactions with growing nascent protein, we developed a model that allowed us to fit the data for multiple chain lengths globally (see Methods and Supplementary Information for details). The time constant of the fastest component (τ_1_) is shared by all constructs and therefore likely corresponds to dissociation of trigger factor from the ribosome without contributions from its interactions with nascent chains. We therefore treated τ_1_ as a global parameter in our analyses. Up to three additional time constants (τ_2_ – τ_4_) are required for all constructs; these are treated as local parameters which are allowed to have different values for every construct. Treating τ_1_ as a global parameter does not significantly affect the quality of the fit, judging by the reduced chi-squared (χ^2^) values. However, adding τ_2_, τ_3_ and τ_4_ to the list of global parameters significantly reduces the fit quality (see Suppl. Table S2 for details). The amplitudes Α_1_ – Α_4_ are also treated as local parameters. The construct RNC32 is adequately described by just three exponentials (Fig. 2 and Suppl. Fig. S4).

Even though the time constants for the three additional components are not linked, they exhibit similar values across all constructs (Fig. 3A and Suppl. Table S3A). However, they contribute to the overall dwell times to different degrees. To quantify their contributions, we calculated the fractional amplitudes of all components (f_1_ to f_4_, Fig. 3B and Suppl. Table S3B). The fastest one (τ_1_) dominates at short chain lengths and accounts for 80% of the dwell time observed for RNC32, consistent with representing direct trigger factor-ribosome interaction. The slowest component (τ_4_) significantly contributes only in constructs with chains lengths of 100 amino acids or more. The contributions of intermediate components (τ_2_ and τ_3_) fluctuate across chain lengths.

**Figure 3.**
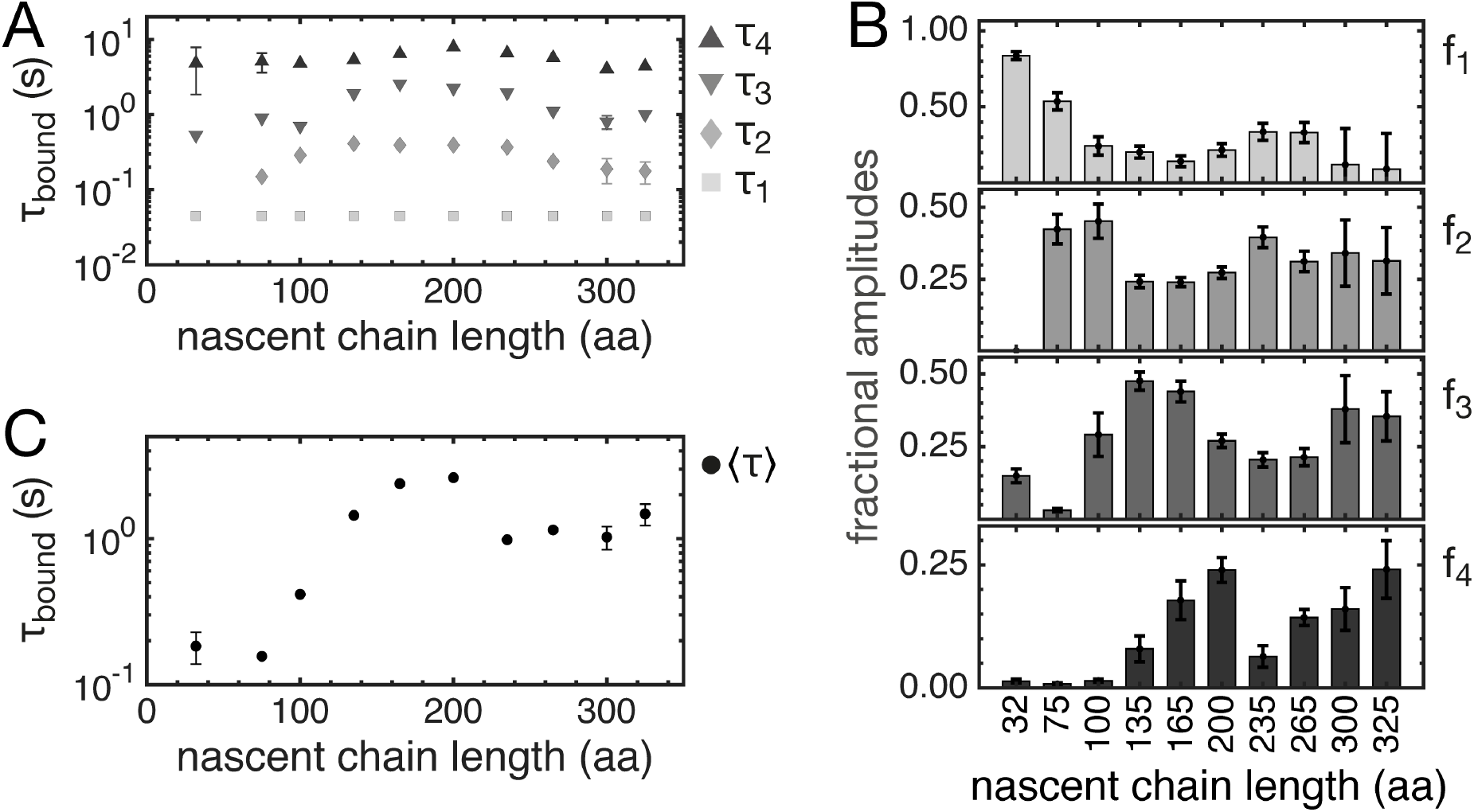
Chaperone-RNC interactions change with increasing nascent chain length. **A**. Time constants obtained from fitting dwell time distributions to multi-exponential models. The fastest time constant (τ_1_) is linked across all datasets as a global fitting parameter. 3 components yield an adequate fit for lengths of L = 32. All other lengths require 4 components. **B**. Fractional amplitudes f for each exponential term shown in A. The fastest component dominates at very short chain lengths, whereas the slower components (τ_2_ to τ_4_) are the main contributors for longer nascent chains. **C**. Ensemble mean dwell times, calculated as the weighted mean of all components. Dwell times start increasing at a length of L = 100 amino acids and peak at L = 200 amino acids. Error bars represent standard deviations in all panels. In A. and C., error bars are not shown if their dimensions on the plot are smaller than the marker size.

Interestingly, the ensemble mean dwell time, <τ>, calculated as the weighted mean from the individual time constants and their fractional amplitudes, does not increase monotonically with nascent chain length, but exhibits a maximum at a length of 200 amino acids (Fig. 3C and Suppl. Table S4). The number of motifs in the nascent protein that can interact with trigger factor should increase with increasing chain length, and they may differ in their affinity for binding sites in the chaperone. In addition, the positioning of interacting motifs relative to binding sites in trigger factor changes as the nascent chain grows in length. The observed change in overall stability thus reflects a complex network of transient interactions that changes during nascent chain elongation.

### Trigger factor binding is destabilized by forced extension of the nascent protein

The substrate binding sites in trigger factor are in close spatial proximity to each other (20) and to the end of the ribosomal polypeptide exit tunnel (29) (Suppl. Fig. S6). Even relatively short nascent polypeptides can occupy multiple binding sites simultaneously if the interaction motifs in the nascent protein are appropriately positioned in space.

Previous work (16) suggested that interacting motifs in the client protein are not adjacent, but distributed across the sequence, resulting in nascent chain compaction upon binding to trigger factor. Increasing the spatial distance between binding sites by applying an external perturbation is thus expected to reduce the overall stability of trigger factor-RNC complexes.

To test this idea experimentally, we combined fluorescence detection with mechanical manipulation using optical tweezers (30). The partially synthesized EF-G G-domain remains unfolded (22, 31) and is therefore expected to be mechanically pliant and extend under an applied external load. Forced stretching is thus expected to disfavor multivalent interactions with the closely spaced binding sites in trigger factor. We tethered single RNC135 molecules between two optically trapped polystyrene beads in the presence of Atto-532-labeled trigger factor (Fig. 4A) and applied force by moving one of the traps relative to the other. Transient spikes in fluorescence emission were detected by confocal fluorescence excitation at the position of the tethered RNC (Fig. 4B), but not in neighboring regions (Suppl. Fig. S5), representing trigger factor binding events from which we extracted dwell times.

**Figure 4.**
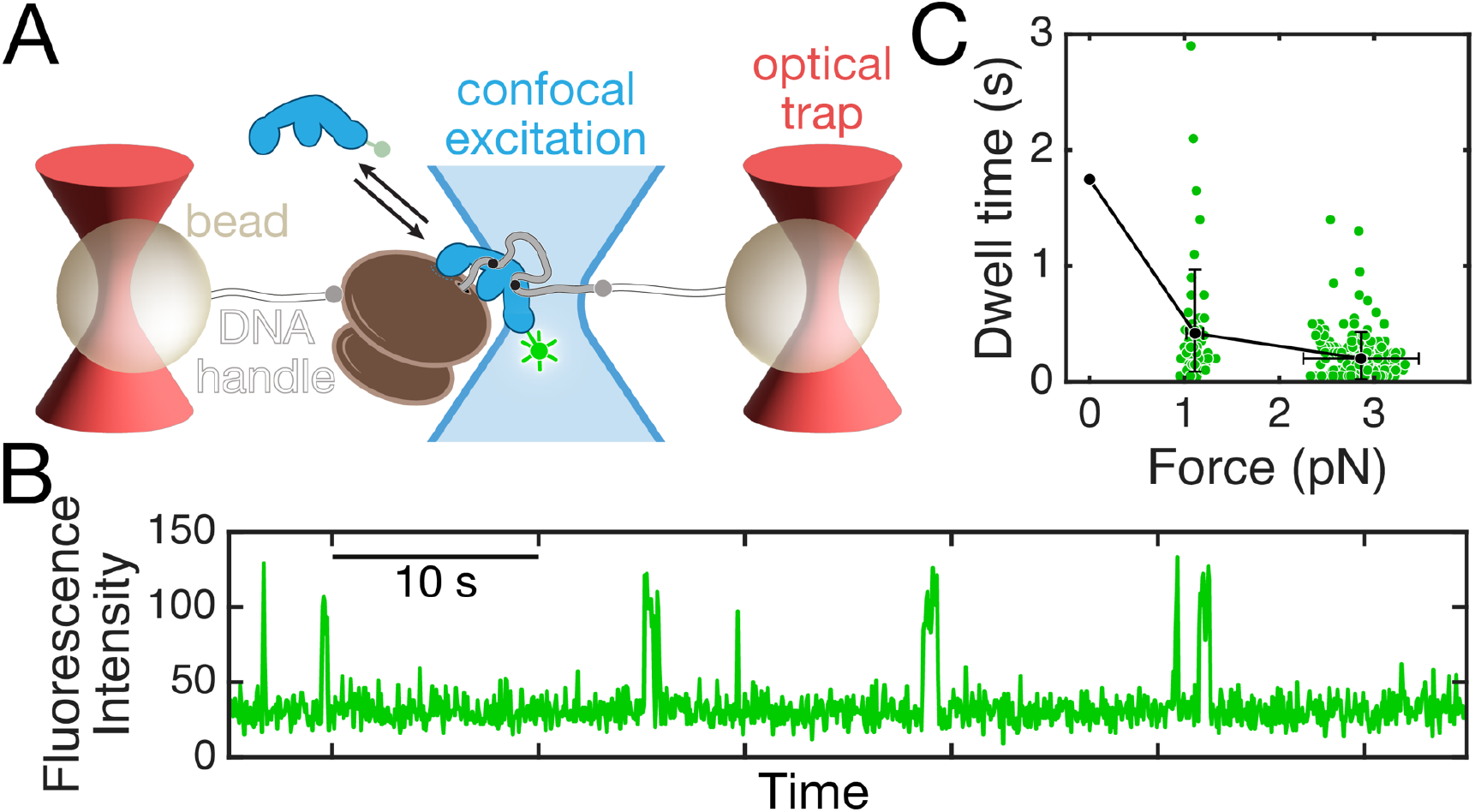
Trigger factor-nascent chain interactions are sensitive to mechanical force. **A**. Schematic of the optical tweezers setup with confocal fluorescence detection (not drawn to scale). RNC135 molecules are tethered between two optically trapped beads by means of long (5 kbp) DNA handles. Moving one of the optical traps enables mechanical manipulation of the RNC, while binding of single trigger factor molecules is detected by confocal fluorescence excitation. **B**. Example fluorescence vs. time trajectory of a point scan recorded at the position of the tethered RNC with a frame rate of 20 Hz. Brief spikes in fluorescence intensity upon excitation at 532 nm correspond to Atto-532 labeled trigger factor binding to the tethered RNC. **C**. Trigger factor dwell times on RNC135 at forces around 1 and 3 pN. Green circles represent individual events, black dots represent the mean. Error bars: s.d. The mean dwell time from TIRF experiments (Fig. 3) is shown at a force of 0 pN for reference.

Holding the nascent chain at forces around 1 pN resulted in a clear shift of dwell times toward lower values, compared to those obtained from TIRF experiments in the absence of force (Suppl. Fig. S5), indicating faster dissociation of the chaperone. Increasing the applied load to approximately 3 pN further decreased the dwell times (Suppl. Fig. S5). Using a single-exponential parameter estimate yielded time constants for dissociation of 0.42 s and 0.20 s at 1.1 and 2.9 pN, respectively (Fig. 4C, Suppl. Fig. S5). Applying an external mechanical load therefore results in a significant acceleration of trigger factor dissociation from RNC135 (Fig. 4C), supporting the idea that a network of dynamic interactions between the chaperone and its unfolded nascent chain client stabilizes binding.

### Nascent chain flexibility shapes chaperone interaction profiles

Trigger factor contains four known nascent chain binding sites that simultaneously interact with binding motifs in suitably long nascent polypeptides, resulting in a complex and changing network of interactions that determine the dwell time measured in TIRF experiments. To conceptualize these findings, we modeled unfolded EF-G chains as worm-like chains (WLC) (32), which allows calculation of the probability distribution in space and, thus, the effective concentration (c_eff_) of any position in the polypeptide relative to another position (33). To this end, we generated discrete WLC models that are restricted to one hemisphere, mimicking the steric exclusion from the ribosome (Fig. 5A, see Supplementary Information for details).

**Figure 5.**
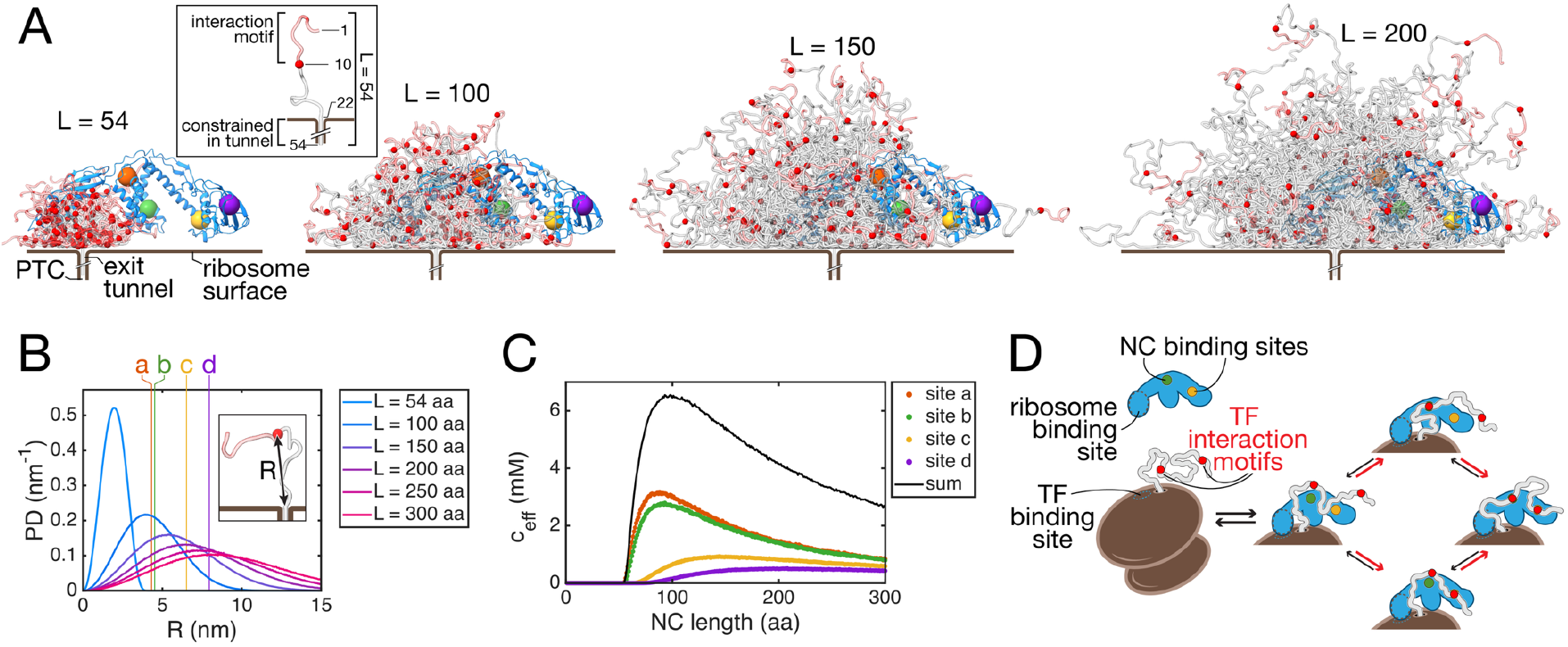
Multi-valent chaperone-RNC contacts change with increasing nascent chain length. **A**. Illustration of unfolded nascent chain conformations. 100 discrete worm-like chain models with the indicated number of amino acids each are shown in cartoon representation. The nascent chain is modeled with 32 C-terminal amino acids sequestered in the exit tunnel of the ribosome and a 10 amino acid binding motif at the N-terminus (inset). Red spheres indicate the position of residue 10. The structure of ribosome-bound trigger factor (pdb code: 1w2b) is shown as a cartoon representation in blue, with colored spheres indicating positions of the nascent chain binding sites ‘a’ – ‘d’. **B**. Probability distribution of distance (R) from the interaction motif to the tunnel exit for nascent chains ranging from 54 to 300 amino acids, calculated numerically from 10^8^ WLC models (illustrated in panel A). Vertical lines indicate the distance of the four substrate binding sites (‘a’ – ‘d’) within trigger factor as described by Saio *et al*. (20). **C**. Effective concentration (c_eff_) of the interaction motif at the positions of binding sites in trigger factor, calculated numerically from the WLC models (see Supplementary Information). **D**. Cartoon illustration of trigger factor interactions with nascent chains. Only two of the four nascent chain binding sites in trigger factor are shown (in green and yellow). Two trigger factor interaction motifs in the nascent chain are indicated by red circles. The two segments can simultaneously bind trigger factor, with on-rates (red arrows) that depend, due to local concentrations, on their positions relative to the ribosome and relative to each other. This binding mode gives rise to complex dissociation kinetics because complete dissociation of the chaperone requires simultaneous dissociation of nascent chain and ribosome contacts.

Considering that approximately 32 amino acids are sequestered within the polypeptide exit tunnel, a nascent chain of 54 amino acids in length is in principle sufficient for an interaction motif to reach the closest binding site in trigger factor, termed site ‘a’ (using the nomenclature from Saio *et al*. (20)), which is 4.3 nm away from the tunnel exit (Suppl. Fig. S6) (see Supplementary Information for details). However, due to the short persistence length of unfolded polypeptides, the probability of finding the interaction motif near this binding site is close to zero (Fig. 5 A,B). At a length of 100 amino acids, the probability of the interaction motif being close to binding site ‘a’ and the nearby site ‘b’ is maximal, and the probability increases for sites ‘c’ and ‘d’ in longer chains (Fig. 5 A,B). For short nascent chains, the stability of the chaperone-RNC complex is therefore dominated by direct interaction with the ribosome, whereas stabilization by nascent chain-interactions only contributes at longer chain lengths. This simplified model explains the increase in stable binding at around 100 amino acids observed by us (this work) and others (17).

## Discussion

Several recent studies have investigated the consequences of TF interactions on nascent client protein stability, conformation and folding using a variety of experimental approaches (14, 16, 26, 34–41) as well as binding of the chaperone to ribosomes and RNCs with nascent chains in cells (17, 28, 31, 42, 43). In contrast, detailed quantitative studies of the interaction dynamics that underpin these observations remain sparse (13, 20). We have developed single-molecule fluorescence approaches to enable direct observation and kinetic characterization of chaperone-nascent chain interaction on the ribosome. Results from these measurements are overall consistent with previously reported values and resolve important details of trigger factor with an authentic nascent chain client.

Trigger factor binds in a dynamic and complex mode to nascent chains of the unfolded G-domain from EF-G. The observed mean dwell times range from 1 to 3 s for nascent chains longer than 100 amino acids, whereas shorter chains are bound for ~200 to 400 ms. Tracking experiments in live cells yielded a global average of ~1 s for the residence time of trigger factor on translating ribosomes (28), in line with the values determined here. Interestingly, trigger factor-RNC complexes exhibit multi-phasic dissociation kinetics, with mean dwell times that increase with chain lengths up to 200 amino acids and then decreases to slightly lower values.

Individual domains found in natural proteins are relatively small, consisting of approximately 100 amino acids on average (44). Stable co-translational folding is therefore possible in this length range for many proteins, changing interactions with chaperones. The choice of EF-G client nascent chains that remain unfolded eliminates the added complexity that results from stable structure formation during folding or misfolding. Our measurements thus provide a detailed, extended view of chaperone-nascent chain interactions before folding (or misfolding) competes with or may be promoted by chaperone binding (16).

Modeling the unfolded nascent protein as a worm-like chain yields the effective concentrations of interaction motifs in the nascent chain at the individual binding sites in trigger factor, calculated from the probability distributions from WLC models. These effective concentrations change as elongation proceeds (Fig. 5C). Interactions with individual motifs are highly dynamic, with K_D_ values in the high micromolar range (20), and thus rearrange on a relatively fast timescale. Binding of one motif to trigger factor constrains the fast conformational rearrangements of the unfolded nascent polypeptide and thus the local concentrations of nearby motifs, which in turn determines the on-rate for another binding event (Fig. 5D). Dwell times are therefore expected to strongly depend on the changing availability of interaction sites during elongation and their positioning relative to the four nascent chain binding sites in trigger factor, which explains the non-monotonic dependence of dwell times on nascent chain lengths observed in TIRF experiments (Fig. 3C).

The simple WLC-behavior can thus explain the multi-phasic dissociation kinetics and peaked mean dwell time distribution that we observe experimentally (Fig. 3C), even though our measurements cannot resolve individual interactions. The observed reduction in dwell times upon mechanical stretching of the nascent protein (Fig. 4) support this model of multivalent binding that allows rapid conformational sampling of the nascent protein without full dissociation from trigger factor. The simple WLC model used here does not take into account interactions of the nascent polypeptide with the surface of the ribosome, which have a profound effect on the conformational space explored by unfolded nascent proteins (14, 45, 46).

Ribosomes translate about 10 – 20 codons per second in exponentially growing *E. coli* cells (47, 48). The ensemble mean dwell time of 2 seconds determined here results in trigger factor dissociation after synthesis of approximately 40 amino acids or fewer. Given an intracellular trigger factor concentration of 50 µM, the chaperone is expected to rapidly rebind to RNCs. However, binding to the nascent protein through a dynamic network of interactions allows the elongating protein to sample conformational space in search of stable structures even while the chaperone is still bound. Notably, while the interaction of trigger factor with its docking site on the ribosome (7) has the shortest dwell time in our kinetic scheme (τ_1_ in Fig. 3), a trigger factor mutant defective in ribosome binding yields very few detected binding events (Fig. 1D). Despite the stabilization from dynamic interactions with the client nascent chain, ribosome binding is thus the decisive step in trigger factor-substrate interaction.

Taken together, our single-molecule experiments developed here open new avenues to explore the mechanisms that support chaperone function at the molecular level. The data suggests a multivalent mode of trigger factor binding to nascent chains that stabilizes the not-yet-folded emerging protein against non-productive interactions while allowing conformational sampling in search of the native state. This constant surveillance throughout synthesis may also guard against misfolding at early stages of protein biogenesis, preempting deleterious downstream consequences.

## Materials and Methods

### Expression, purification and labeling of trigger factor

DNA amplicon encoding wild type *E. coli* trigger factor sequence with a single cysteine mutation (T150C) was cloned into a pBAD-His_6_ tag-SUMO vector with a Ulp1 cleavage site (49) and an ampicillin resistance cassette. The mutant protein defective in ribosome binding had positions 44-46 replaced with Alanine (FRK > AAA) (7). Plasmids were transformed into chemically competent *E. coli* BL21 strain cells and grown overnight on an agar plate with carbenicillin (Thermo Scientific). Colonies grown overnight were scraped together and inoculated into LB growth media along with carbenicillin and the culture was grown at 37°C to an OD_600_ of ~0.5-0.6. The protein expression was induced by the addition of 0.2% (w/v) L-arabinose, and the culture was grown for 3 hours at 37°C. The cells were pelleted down by centrifuging at 5000 rpm for 30 min at 4°C, washed with lysis buffer (50 mM HEPES-KOH at pH 7.4, 150 mM KCl, 5% glycerol), pelleted again and the cell pellets were stored at −80°C. Wild-type and mutant trigger factor was purified using metal affinity chromatography, followed by proteolytic removal of the His_6_ tag and size exclusion chromatography (see Supplementary Information for details). For fluorescence experiments, trigger factor (wild-type and FRK/AAA mutant) was labeled at the engineered cysteine residue at position 150 using Atto-532 maleimide and purified by size exclusion chromatography. Details are described in Supplementary Information.

### Preparation and fluorescent labeling of ribosome-nascent chain complexes (RNCs) for TIRF experiments

RNCs for TIRF microscopy experiments were generated by in vitro methods. DNA templates for the non-stop G-domain constructs with a T7 promoter, ribosome binding site, and a SpyTag were generated by PCR. DNA templates were used to synthesize mRNAs by *in vitro* transcription (Megascript T7 transcription kit, Invitrogen). Biotinylated oligos were annealed to the 5’-end of the mRNAs by denaturing the mRNA and slowly cooling the mixture of oligos and mRNA. mRNAs annealed to oligos were used to synthesize stalled RNCs by *in vitro* translation (PURExpress *in vitro* protein synthesis kit, New England Biolabs).

SpyCatcher with a N-terminal cysteine (27) was labeled with maleimide-modified sulfo-Cy5 (Lumiprobe) similar to the procedure used for labeling trigger factor. Labeled and purified SpyCatcher-sulfoCy5 was mixed with stalled RNCs after in vitro translation and incubated at room temperature for 30 minutes. SpyCatcher forms a covalent isopeptide bond with SpyTag on the nascent chain. Sulfo-Cy5 label on the nascent chain enable the identification of the RNCs in TIRF experiments. RNC mixture was pelleted by ultracentrifugation to remove excess SpyCatcher-sulfoCy5 and other translation components. RNC pellet was resuspended in storage buffer (20 mM HEPES, pH 7.4, 100 mM KCl, 20 mM MgCl_2_, 5 mM β-mercaptoethanol, 1-2 units of RNaseOUT recombinant ribonuclease inhibitor (Invitrogen)) and flash frozen. The molecular weight and the concentration of RNCs were estimated by bis-tris polyacrylamide gel electrophoresis.

### Preparation of ribosome-only control

To prepare RNC_control_ (without any EF-G sequence), a shorter mRNA consisting of 25 amino acids was prepared by in vitro transcription and annealed to the biotinylated oligo. A nascent chain consisting of 25 amino acids is expected to be completely buried inside the ribosomal exit tunnel. Ribosomes containing C-terminally SpyTagged ribosomal protein L17 were used for in vitro translation of the short mRNA and SpyCatcher-sulfoCy5 was used to label the ribosomes directly. Sulfo-Cy5 label on the ribosomes enable the identification of the RNCs in TIRF experiments.

### Preparation of RNCs for optical tweezers experiments

Non-stop mRNAs containing T7 promoter, ribosome binding site, N-terminal SpyTag and the first 135 residues of elongation factor G were generated by in vitro transcription similar to the procedures described above. Ribosomes containing C-terminally SpyTagged ribosomal protein L17 were used for in vitro translation to generate stalled RNCs. After translation, stalled RNCs were pelleted down to remove translation factors, the pellet was resuspended in storage buffer (20 mM HEPES, pH 7.4, 100 mM KCl, 20 mM MgCl_2_, 5 mM β-mercaptoethanol) and flash frozen.

### Single-molecule TIRF microscopy setup and RNC-TF colocalization experiments

Single-molecule fluorescence measurements were performed on a prism-type total internal reflection fluorescence (TIRF) microscope. Spot detection and background subtraction were carried out following established protocols (50). Data acquisition was controlled using smCamera software (51–53) with a time resolution of 100 ms per frame. Data processing yielded time traces of fluorescence intensities that were used for further analyses. See Supplementary Information for details.

### Data processing for single-molecule TIRF experiments

Red and green emission intensities corresponding to individual molecular time traces were visualized using MATLAB scripts. The red intensities from the first 10 frames corresponding to the excitation of the RNCs were added together and molecules with very low and high intensities (10-15% on either end) were discarded from further analysis. The green intensities from the selected molecules were visualized using MATLAB scripts and molecules with high background were discarded. The remaining molecules were used for estimation of dwell times.

A model-free machine learning approach, DISC (54) was used to identify the low- and high-intensity regions in the green time traces. An alpha-value of 0.05 was used for change point detection, BIC-GMM was used for both divisive and agglomerative clustering and 1 iteration of the Viterbi algorithm was used for identifying the high- and low-intensity states. The number of states was fixed at 2 to group all high-intensity states together. The low- and high-intensity states are referred to as unbound and bound states in the manuscript respectively. The assigned states were converted to time intervals and data from multiple movies from different days were grouped together for analysis.

### Optical tweezers experiments

Ribosomes with a C-terminal SpyTag on the ribosomal protein L17 were used to prepare RNCs with a N-terminal SpyTag on the nascent chain following published procedures (27). All single-molecule optical tweezers experiments were performed using a commercial instrument (C-Trap, Lumicks). Fluorescence co-localization experiments were carried out in the presence of 2 nM Atto-532 labeled trigger factor. See Supplementary Information for details.

### Extracting kinetic parameters from TIRF data

To obtain kinetic parameters from dwell times measured in TIRF experiments, we fit a model function to binned dwell times. The model function was the difference between the values of the survival function *S*(*t*) for the *t* values corresponding to the start and the end of each bin. The survival function *S*(*t*) was modeled as a linear combination of a number of exponential functions. The fitting parameters included the absolute amplitudes *Α*_*i*_ and time constants *τ*_*i*_ associated with each exponential term (see Supplementary Information for details). The fitting was performed by a weighted least squares estimator. The output of the program included all *Α*_*i*_ and *τ*_*i*_ and also the fractional amplitudes *f*_*i*_. A standard deviation was calculated by the program for each output value.

### Estimation of dwell times from optical tweezers experiments

Green fluorescence emission intensities were processed at a frame rate of 20 Hz, and the states were assigned using a model-free machine learning approach DISC described above. An alpha-value of 0.05 was used for change point detection, BIC-GMM was used for both divisive and agglomerative clustering and 1 iteration of the Viterbi algorithm was used for identifying the high- and low-intensity states. The number of states was fixed at 2 to group all high-intensity states together. The low- and high-intensity states are referred to as unbound and bound states in the manuscript respectively. A force baseline collected in the absence of a molecular tether was subtracted from the force recordings and plotted against the binding dwell times. A total of 292 events from 12 individual molecules were used for the analysis. The mean dwell time was obtained from an exponential parameter estimate using the Matlab function expfit.

### Worm-like chain models and calculation of effective concentrations

To model the interactions with trigger factor binding sites, we modeled the nascent chain as containing three elements: (1) a 32 amino acid segment confined within the ribosomal polypeptide exit tunnel, (2) a flexible region modeled as a worm-like chain (WLC), and (3) a trigger factor interaction motif of 10 amino acids (the average motif length reported previously (20). Parameters for the WLC model were a persistence length of 0.50 nm and a contour length of 0.36 nm per amino acid. WLC simulations were restricted to the z>0 half-space, assuming that the z≤0 half-space was occupied by the ribosome. Residue 41 (from the PTC) is the first residue outside the tunnel with an initial direction normal to the ribosome surface. Subsequent random changes in the direction of the WLC were simulated as described in Supplementary Information. For each chain length, 10^8^ random WLCs were simulated. Distances (R) of the interaction motif from the tunnel exit to the free end of the nascent chain were calculated from the simulated models and binned using a bin width of 0.04 nm. The concentration of the interaction motif at the locations of binding sites in trigger factor were calculated from the simulated models assuming a maximum distance of 0.2 nm for each of the binding site (a, b, c and d) identified by Saio *et al*. (20), using coordinates from the structure of ribosome-bound trigger factor (29).

## Acknowledgments

This work was supported by a Grant from the National Institutes of Health (5R01GM121567) to C.M.K./V.J.H. The work in the Ha lab was funded by a grant from the NIH (R35 GM 122569).

