## Supplementary Information for "Multivalent weak contacts shape chaperone-nascent protein interactions"

This file contains

- Supplementary Figures
    - Fig. S1....2
    - Fig. S2....3
    - Fig. S3....4
    - Fig. S4....5
    - Fig. S5....6
    - Fig. S6....7
- Supplementary Tables
    - Table S1....8
    - Table S2....9
    - Table S3A....10
    - Table S3B....11
    - Table S4....12
- Supplementary Methods
    - Purification of trigger factor....13
    - Fluorescent labeling of trigger factor....13
    - Single-molecule TIRF microscropy....14
    - Single-molecule RNC-TF colocalization experiments....14
    - Optical tweezers experiments....14
    - Dwell time data analysis....15
    - Suppression of systematic errors....18
    - Worm-like chain simulation procedure....18

#### Supplementary Figures

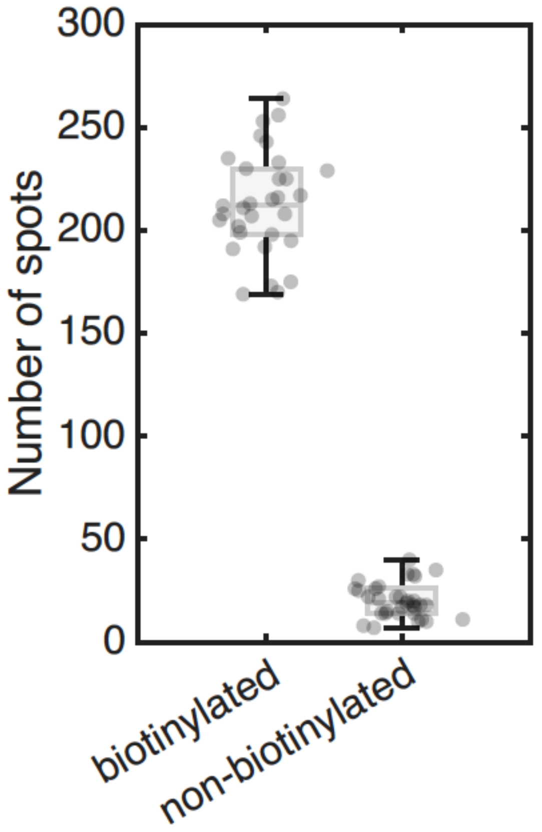


**Suppl. Fig. S1: Labeled RNCs are specifically attached to the surface.** The number of red spots observed after excitation with a 638-nm laser for 10 frames is shown for two constructs: RNC 200 prepared from a mRNA annealed to a biotinylated oligo and RNC 135 prepared from a mRNA annealed to a non-biotinylated oligo. The number of spots from multiple short movies is represented as individual markers offset horizontally for clarity (n = 33 for biotinylated construct and n = 31 for non-biotinylated construct). The distribution of the data is represented as a box and whisker plot, with the box showing the interquartile range and the median line. The experiments for the biotinylated and non-biotinylated constructs were performed on the same day to enable a direct comparison.

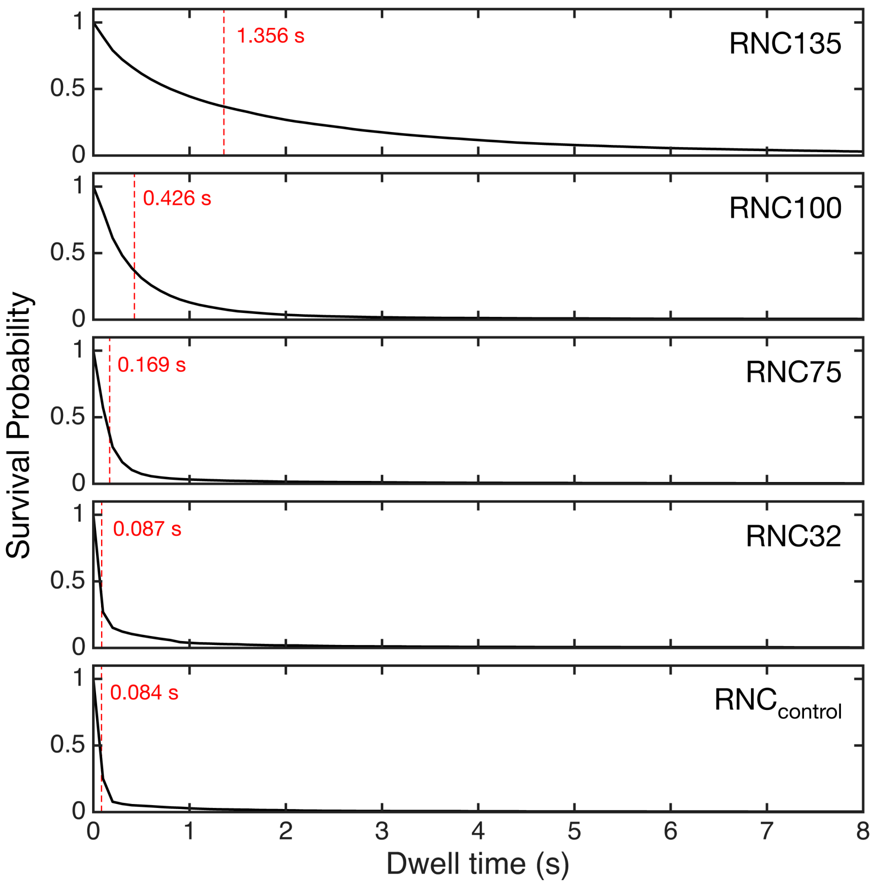


**Suppl. Fig. S2. Experimental survival functions for trigger factor bound to RNCs.** The survival function $S(t)$ represents the probability that the dwell time is greater than $t$ and is calculated as a ratio of the number of dwell times greater than $t$ to the number of all registered dwell times. A dashed red line in each panel represents $t_{1/e}$, which is defined such that $S(t_{1/e})=1/e$; for an exponential survival function $t_{1/e}$ would be equal to the mean dwell time.

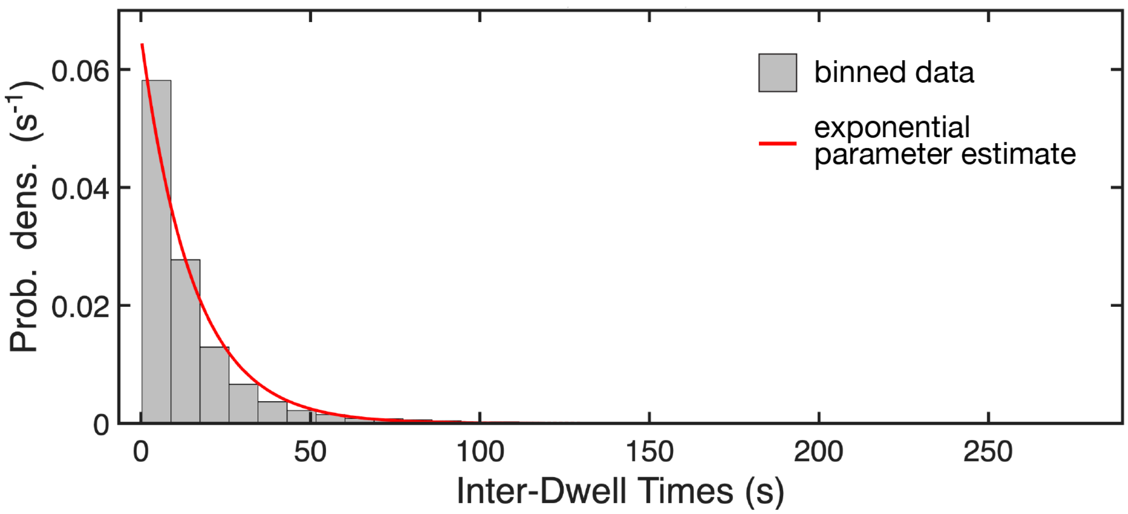


**Suppl. Fig. S3. Analysis of inter-dwell times to estimate association rate constants.** Grey bars represent the distribution of time intervals between binding events, the red line is a single-exponential parameter estimate based on a maximum likelihood estimator.

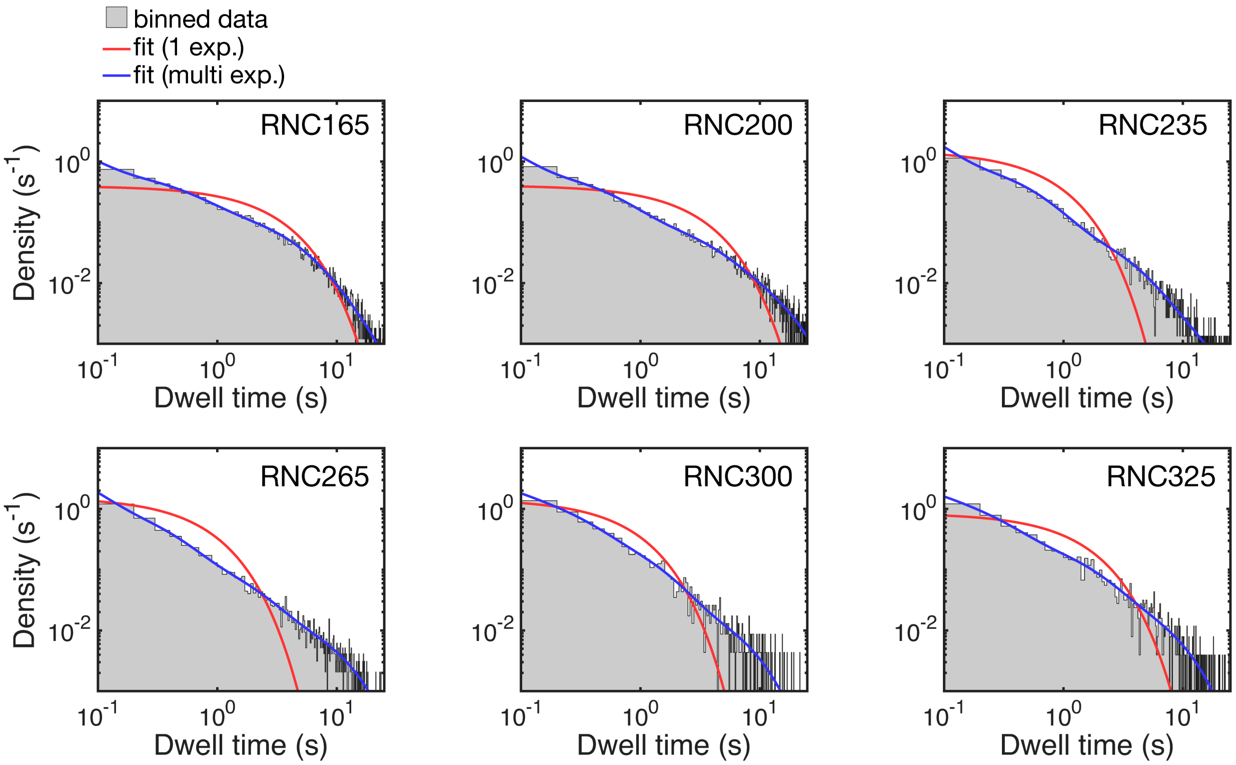


**Suppl. Fig. S4. Dwell time distribution and fits for trigger factor binding to RNCs with nascent chains of varying lengths.** Binned dwell times and fits for RNCs with chain lengths above 150 amino acids. A single-exponential model (red line) does not describe the binned dwell times (grey area) well, whereas adequate fits are obtained with a multi-exponential model (blue line) containing 4 terms.

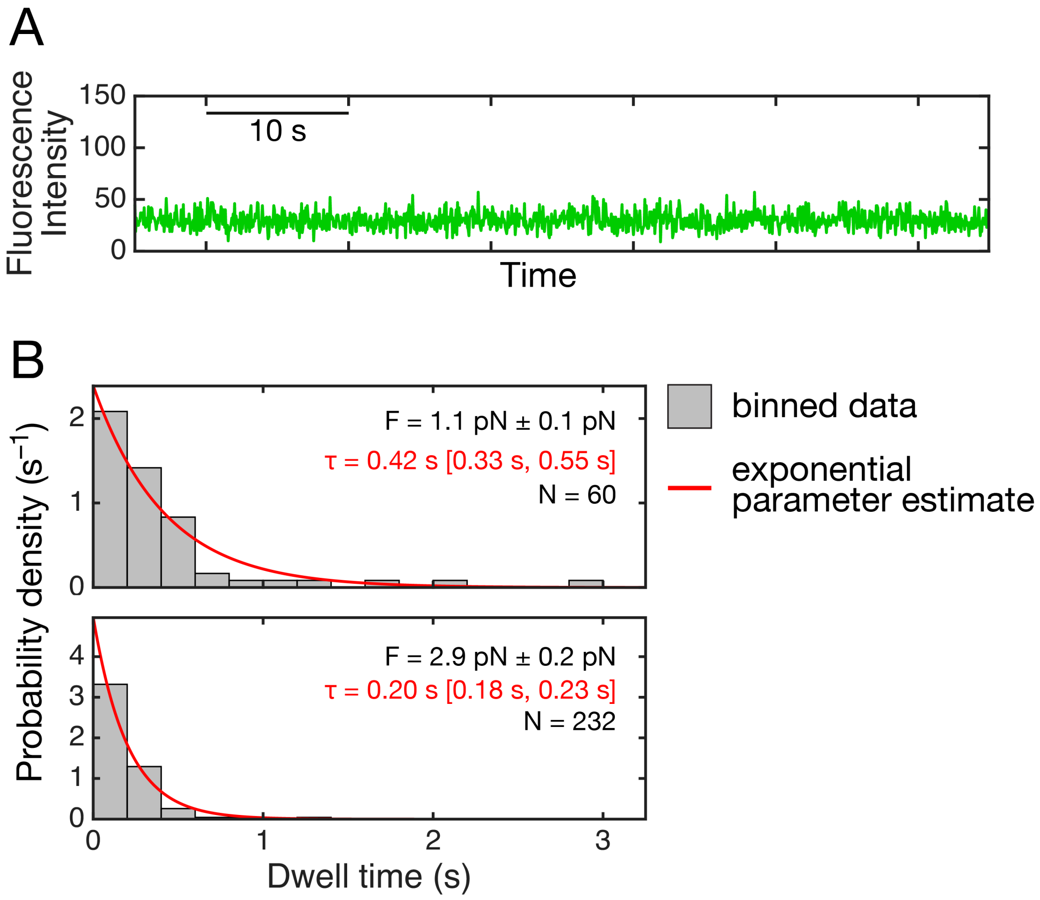


**Suppl. Fig. S5. A.** Example trajectory of a point scan recorded in a control region away from trapped beads during the same experiment as shown in Fig. 4B in the main text. A 532-nm laser excitation was used and the fluorescence intensity at a single coordinate was recorded at a frequency of 20 Hz. **B.** Dwell time distributions for trigger factor binding to RNC135 under mechanical load. Grey bars represent binned data of dwell times measure at force of approximate 1 and 3 pN, red lines represent single-exponential parameter estimates. Comparing the two distributions using a two-sample Komolgorov-Smirnov test yielded a p-value of 3.7, indicating that the distributions differ in a statistically significant way. Force values are provided as means ± standard deviation, time constants (τ) as mean and 95% confidence intervals.

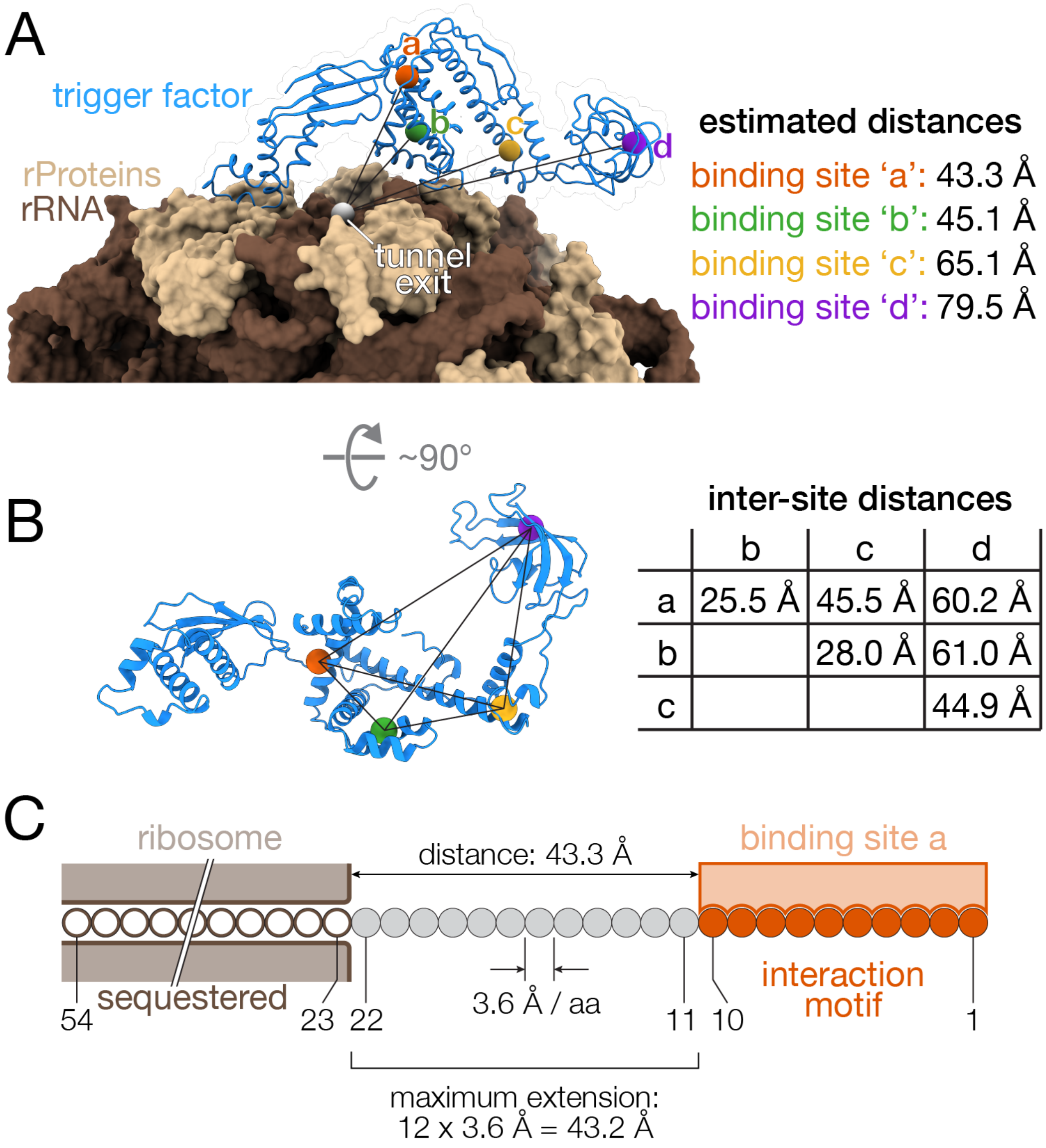


**Suppl. Fig. S6.** **A.** Distances of nascent chain binding sites 'a' to 'd' in trigger factor relative to the tunnel exit. **B.** Distances of nascent chain binding sites 'a' to 'd' relative to each other. **C.** Diagram illustrating the model used for estimating probability densities and local concentrations of interaction motifs at trigger factor binding sites (Fig. 5 in the main text). The diagram illustrates the minimal length needed for an N-terminal interaction motif (residues 1 – 10) to reach the closest nascent chain binding site (site 'a') in ribosome-bound trigger factor. Distances in panels A and B were estimated from pdb structures 1w26/1w2b.

#### Supplementary Tables

| | $N_{exp}$ | $RNC_{control}$ | RNC32 | RNC75 | RNC100 | RNC135 |
|---|---|---|---|---|---|---|
| reduced $\chi^2$ | 1 | 5.162 | 2.623 | 3.948 | 2.697 | 4.733 |
| | 2 | 1.024 | 1.385 | 1.556 | 1.343 | 1.409 |
| | 3 | 0.984 | 1.198 | 1.125 | 0.994 | 1.098 |
| | 4 | N. C. | N. C. | 0.992 | 0.966 | 1.026 |
| | 5 | N. C. | N. C. | N. C. | 0.965 | 1.032 |
| AIC | 1 | 330.8 | 175.6 | 441.3 | 395.7 | 1084.6 |
| | 2 | 5.9 | 44.4 | 85.7 | 82.9 | 121.8 |
| | 3 | 4.8 | 26.4 | 24.1 | 4.7 | 33.9 |
| | 4 | N. C. | N. C. | 6.8 | 0.4 | 15.2 |
| | 5 | N. C. | N. C. | N. C. | 2.1 | 19.1 |

**Suppl. Table S1: Reduced $\chi^2$ and AIC for the fits of multi-exponential model functions to dwell time data.** Single-set (not global) fits to binned experimental dwell times were characterized by the reduced $\chi^2$ and by the Akaike Information Criterion (AIC). The results are shown for five constructs with nascent chain lengths between 0 and 135 aa. The fits were obtained using the models with 1, 2, 3, 4, and 5 exponential terms. In those cases where the data set contained insufficient information to unambiguously estimate all the fitting parameters, the least square estimator algorithm did not converge; the corresponding cells are marked "N. C.". The lowest value of AIC for each construct is underlined; it indicates the optimum number of exponential terms for that construct.

| global parameters | $N_D$ | $N_P$ | $\chi^2$ | reduced $\chi^2$ | AIC |
|---|---|---|---|---|---|
| none | 2904 | 82 | 2941.7 | 1.042 | 201.7 |
| $\tau_1$ | 2904 | 72 | 2968.6 | 1.048 | 208.6 |
| $\tau_1$, $\tau_2$ | 2904 | 62 | 3187.0 | 1.121 | 407.0 |
| $\tau_1$, $\tau_2$, $\tau_3$ | 2904 | 53 | 3225.2 | 1.131 | 427.2 |
| $\tau_1$, $\tau_2$, $\tau_3$, $\tau_4$ | 2904 | 45 | 3275.8 | 1.146 | 461.8 |

**Suppl. Table S2: Statistical characteristics of global fits to all data sets by four-exponential model function.** $N_D$ is total number of bins with non-zero weights in all data sets. $N_P$ is the total number of all global and local fitting parameters (this number takes into account the fact that only two exponential terms were used for $RNC_{control}$ and three exponential terms were used for RNC32). The $\chi^2$ value was calculated using Suppl. Equation (7). Reduced $\chi^2$ was calculated as $\chi^2/(N_D$-$N_P)$. The Akaike Information Criterion (AIC) was calculated as $2N_P+\chi^2-N_D$.

| Construct | $\tau_1$ (s) | $\sigma\tau_1$ (s) | $\tau_2$ (s) | $\sigma\tau_2$ (s) | $\tau_3$ (s) | $\sigma\tau_3$ (s) | $\tau_4$ (s) | $\sigma\tau_4$ (s) |
|---|---|---|---|---|---|---|---|---|
| $RNC_{control}$ | 0.0449 | 0.0039 | 1.0132 | 0.1183 | | | | |
| RNC32 | 0.0449 | 0.0039 | 0.5325 | 0.0438 | 4.8506 | 2.9965 | | |
| RNC75 | 0.0449 | 0.0039 | 0.1492 | 0.0078 | 0.9007 | 0.1550 | 5.1046 | 1.4830 |
| RNC100 | 0.0449 | 0.0039 | 0.2876 | 0.0330 | 0.7027 | 0.0711 | 4.8249 | 0.8618 |
| RNC135 | 0.0449 | 0.0039 | 0.4106 | 0.0440 | 1.9159 | 0.1339 | 5.3428 | 0.7273 |
| RNC165 | 0.0449 | 0.0039 | 0.3920 | 0.0331 | 2.5656 | 0.1988 | 6.4714 | 0.5392 |
| RNC200 | 0.0449 | 0.0039 | 0.3924 | 0.0331 | 2.2523 | 0.2475 | 7.8922 | 0.3942 |
| RNC235 | 0.0449 | 0.0039 | 0.3676 | 0.0278 | 1.9620 | 0.2915 | 6.6016 | 1.2174 |
| RNC265 | 0.0449 | 0.0039 | 0.2392 | 0.0291 | 1.1164 | 0.1413 | 5.7239 | 0.3326 |
| RNC300 | 0.0449 | 0.0039 | 0.1889 | 0.0693 | 0.8029 | 0.1609 | 4.0586 | 0.5537 |
| RNC325 | 0.0449 | 0.0039 | 0.1765 | 0.0577 | 1.0052 | 0.1879 | 4.4108 | 0.4410 |

**Suppl. Table S3A: Detailed fit results.** Time constants $\tau_i$ and standard deviations $\sigma\tau_i$ associated with the four exponential terms in the survival function, see Suppl. Equation (1).

| Construct | $f_1$ | $\sigma f_1$ | $f_2$ | $\sigma f_2$ | $f_3$ | $\sigma f_3$ | $f_4$ | $\sigma f_4$ |
|---|---|---|---|---|---|---|---|---|
| $RNC_{control}$ | 0.9615 | 0.0069 | 0.0385 | 0.0069 | | | | |
| RNC32 | 0.8366 | 0.0258 | 0.1497 | 0.0239 | 0.0136 | 0.0039 | | |
| RNC75 | 0.5365 | 0.0567 | 0.4242 | 0.0518 | 0.0313 | 0.0056 | 0.0080 | 0.0023 |
| RNC100 | 0.2423 | 0.0596 | 0.4519 | 0.0591 | 0.2913 | 0.0749 | 0.0145 | 0.0033 |
| RNC135 | 0.2026 | 0.0396 | 0.2422 | 0.0217 | 0.4760 | 0.0311 | 0.0792 | 0.0265 |
| RNC165 | 0.1421 | 0.0361 | 0.2401 | 0.0158 | 0.4396 | 0.0356 | 0.1781 | 0.0393 |
| RNC200 | 0.2165 | 0.0417 | 0.2734 | 0.0199 | 0.2704 | 0.0232 | 0.2397 | 0.0252 |
| RNC235 | 0.3354 | 0.0556 | 0.3960 | 0.0357 | 0.2049 | 0.0250 | 0.0637 | 0.0218 |
| RNC265 | 0.3311 | 0.0661 | 0.3121 | 0.0356 | 0.2137 | 0.0295 | 0.1431 | 0.0166 |
| RNC300 | 0.1196 | 0.2382 | 0.3410 | 0.1149 | 0.3790 | 0.1154 | 0.1604 | 0.0438 |
| RNC325 | 0.0908 | 0.2341 | 0.3140 | 0.1152 | 0.3543 | 0.0842 | 0.2409 | 0.0588 |

**Suppl. Table S3B: Detailed fit results.** Fractional amplitides $f_i$ and standard deviations $\sigma f_i$ associated with the four exponential terms in the survival function. Fractional amplitudes were calculated from the absolute amplitudes $A_i$ using Suppl. Equation (10); standard deviations were calculated using Suppl. Equation (9).

| Construct | Number of events | $\langle\tau\rangle$ (s) | $\sigma\langle\tau\rangle$ (s) |
|---|---|---|---|
| $RNC_{control}$ | 2277 | 0.08 | 0.01 |
| RNC32 | 2892 | 0.18 | 0.05 |
| RNC75 | 10811 | 0.16 | 0.01 |
| RNC100 | 11379 | 0.42 | 0.03 |
| RNC135 | 12976 | 1.44 | 0.07 |
| RNC165 | 14813 | 2.38 | 0.09 |
| RNC200 | 10712 | 2.62 | 0.13 |
| RNC235 | 5804 | 0.98 | 0.08 |
| RNC265 | 7692 | 1.15 | 0.09 |
| RNC300 | 2142 | 1.03 | 0.18 |
| RNC325 | 2207 | 1.48 | 0.25 |

**Suppl. Table S4: Mean dwell times for RNC constructs.** Mean dwell times $\langle\tau\rangle$ with standard deviations $\sigma\langle\tau\rangle$ for the RNC constructs. Mean dwell time was calculated using Suppl. Equation (11); standard deviations were calculated using Suppl. Equation (9).

### Supplementary Methods

#### Purification of trigger factor

To purify wild-type and FRAK/AAA mutant trigger factor, cells were resuspended in lysis buffer supplemented with protease inhibitors (cOmplete™, Mini, EDTA-free protease inhibitor cocktail, Roche). Resuspended cells were lysed in a high-pressure homogenizer (C5-emulsiflex, Avestin, Canada) chilled on ice. The cell debris was removed by centrifugation at 40,000 g for 45 minutes at 4 °C and discarding the pellet. Clarified cell lysate was further filtered by a 0.2 µm syringe filter, loaded onto a superloop and injected into a Ni-NTA column (HisTrap, Cytiva). The column was run on an ÄKTA Pure™ protein purification system (Cytiva). Unbound proteins and other biomolecules were washed off by flowing in lysis buffer. The bound proteins were eluted with a gradient of elution buffer containing imidazole (50 mM HEPES-KOH at pH 7.4, 150 mM KCl, 500 mM imidazole, 5% glycerol). Fractions containing the protein were identified by denaturing SDS-PAGE and pooled together. To cleave the N-terminal $His_6$-SUMO tag, $His_6$-tagged Ulp1 protease and DTT were added to the protein and dialyzed against lysis buffer to remove excess imidazole. The mixture of cleaved and uncleaved protein were loaded onto a Ni-NTA column and the cleaved fractions were eluted with lysis buffer. The protein fractions were further purified by size-exclusion chromatography (HiPrep 16/60 Sephacryl S-300 HR column, Cytiva). Fractions containing the purified protein were concentrated using Amicon Ultra centrifugal filters (Millipore Sigma), aliquoted and flash frozen for later use. The concentration of the purified proteins were measured using a NanoDrop™ UV-Vis spectrophotometer (ThermoFisher Scientific) using an extinction coefficient of 17420 $M^{-1}$ $cm^{-1}$.

#### Fluorescent labeling of trigger factor

For labeling of trigger factor constructs with Atto-532 at the engineered cysteine residue (C150), the protein was mixed with TCEP at room temperature for 30 minutes to reduce any disulfide bonds followed by removal of TCEP using Amicon Ultra centrifugal filters (Millipore Sigma). Trigger factor protein was labeled with maleimide-modified Atto-532 dye (ATTO-TEC GmbH, Germany). The dye was first dissolved in DMSO and added to the protein for a final dye concentration of ~1.6 mM and a dye to protein ratio of 25:1. The protein and dye mixture was incubated in the dark at room temperature for 3 hours. The labeling reaction was quenched with the addition of 10 mM β-mercaptoethanol. The protein and dye mixture was centrifuged to remove any aggregates and separated using size exclusion chromatography (Superdex 200 Increase 10/300 column, Cytiva) to remove excess unreacted dye.

The concentration of the unlabeled and Atto-532-labeled proteins were measured at 280 nm and 532 nm respectively using a NanoDrop™ UV-Vis spectrophotometer (ThermoFisher Scientific), with extinction coefficients of 17420 $M^{-1}$ $cm^{-1}$ and 115000 $M^{-1}$ $cm^{-1}$ respectively. The total protein concentration was obtained by using the relation (A280 - A532*correction factor)/(Extinction coefficient of protein) with a correction factor of 0.09, recommended by the manufacturer. The concentration of the labeled protein and total protein was used to estimate the labeling efficiency. The observed labeling efficiency was ~100% which may include errors due to uncertainties in the extinction coefficient of the dye in aqueous solutions.

**Single-molecule TIRF microscropy**

Atto532 and sulfo-Cy5 fluorophores were excited using a 532-nm laser (Coherent Compass 315M) and a 638-nm laser (Cobolt 06-MLD), respectively. Fluorescence signals were collected through a 60x water-immersion objective (Olympus, NA 1.2) and projected onto a back-illuminated electron-multiplying charge-coupled device (EMCCD) camera (iXON, Andor Technology) configured in a dual-view imaging system. In the emission pathway, a long-pass filter (Semrock BLP02-561R-25) was used to suppress residual 532-nm excitation light, and a notch filter (Chroma ZET633TopNotch) was applied to remove 638-nm laser scattering. Donor and acceptor emission channels were spatially separated using a long-pass dichroic mirror (Semrock FF640-FDi01-25X36) within the dual-view optical assembly. PEG-passivated quartz slides and coverslips were assembled into flow chambers as described previously.

**Single-molecule RNC-TF colocalization experiments**

For single-molecule TIRF assays, the biotin-functionalized flow cells were first incubated with neutravidin to sparsely coat the surface, followed by a wash step to remove excess molecules. Biotinylated RNCs were then introduced at a concentration of ~100–500 pM for surface immobilization, and unbound RNCs were subsequently washed out. All experiments were performed in an imaging buffer consisting of 20 mM HEPES (pH 7.4), 100 mM KCl, 5 mM $MgCl_2$, 5 mM β-mercaptoethanol, 1% (w/v) dextrose, 10 units of glucose oxidase from Aspergillus niger (Sigma-Aldrich), 100 units of catalase from Corynebacterium glutamicum (Sigma-Aldrich), and 2 mM Trolox.

Initially, short movies (3 s; 30 frames) were recorded under 638-nm excitation (targeting the sulfo-Cy5 dye) to assess the surface coverage of immobilized RNCs. Next, Atto-532-labeled trigger factor was flowed in at a final concentration of 2.5 nM in the imaging buffer. To observe colocalization, long movies (2000–3000 frames; 3–5 min) were recorded: an initial 10-frame excitation with the 638-nm laser was used to accurately mark the positions of the RNCs, followed by prolonged excitation with the 532-nm laser to monitor the trigger factor binding dynamics. Finally, the extracted signals were converted into molecular time traces for downstream analysis.

**Optical tweezers experiments**

Doubly tagged RNCs were attached to DNA molecular handles for optical tweezers experiments. To prepare the DNA handles, ~200 bp dsDNA with an overhang was labeled with biotin- or digoxigenin-dUTP at multiple positions. Biotin/digoxigenin labeled dsDNA were ligated to ~4.5 kb long restriction enzyme digested dsDNA with complementary overhangs which was later ligated to SpyCatcher-dsDNA. An equimolar mixture of biotin/digoxigenin labeled DNA handles were covalently linked to SpyTags on the ribosome and nascent chain.

Imaging buffer consisted of 20 mM HEPES-KOH, pH 7.4, 100 mM KCl, 5 mM MgCl2, 5 mM β-mercaptoethanol, 2.5 mM protocatechuic acid (Sigma-Aldrich), 50 nM protocatechuate 3,4-dioxygenase (Sigma-Aldrich), and 2 mM Trolox. RNC aliquots were mixed with biotin- and digoxigenin- labeled DNA handles and incubated on ice overnight to form a covalent link between SpyTag and SpyCatcher. Before the experiment, DNA handle-attached RNCs were mixed with anti-digoxigenin cotated 2.1 um polystyrene beads and flown into the microfluidic flow cell. Streptavidin coated 2.1 um polystyrene beads were flown into a different channel. To form a molecular tether, an anti-digoxigenin coated bead and a streptavidin coated bead were trapped in individual optical traps and brought together. Successful formation of a single molecular tether was confirmed by a rise in force at the expected inter-bead distance (~2.5-2.7 um for the DNA handles used in the experiments).

To monitor interactions of trigger factor with tethered RNCs, 2 nM Atto-532 labeled trigger factor was flowed in to monitor the binding to RNCs. To obtain the coordinates of the beads and the tethered molecule, 2-D scans and 1-D line scans were performed after excitation with a 532-nm laser. Brief pulses of green emission between the beads correspond to trigger factor binding to RNCs. To obtain the lifetime of binding events at a higher time resolution, a confocal point scan was performed at the pixel corresponding to fluorescent signal in 2-D line scans. Fluorescence intensities and the corresponding forces at both the beads were recorded at 78125 Hz.

**Dwell Time Data Analysis**

To mathematically model the set of experimentally observed dwell times we utilized the survival function $S(t)$ that predicts how many of the experimentally detected dwell times are expected to be longer than $t$; to distinguish between this survival function and the normalized survival function that appears in Suppl. Fig. S2, we will call it the absolute survival function. The normalized survival function equals the absolute survival function divided by the value of the latter at $t$=0. The absolute survival function was modeled using a linear combination of exponential functions,

$$S(t)=\sum_{i=1}^{N_{\exp}} A_i \exp(-t/\tau_i) \quad (1)$$

where $N_{\exp}$ is the number of exponential terms, $A_i$ are the amplitudes, and $\tau_i$ are the time constants.

The experimental dwell times were binned. The bins were defined in the following manner: bin number 1 began at $t$=0 and ended at $t=t_1$; Bin number $n$ began at $t=t_{n-1}$ and ended at $t=t_n$. Due to the nature of the technique, all the measured dwell times were integer multiples of the video frame time $\Delta t$ = 100 ms, which compelled us to use bins of the width $\Delta t$, resulting in $t_n=n\Delta t$. All dwell times equal to $n\Delta t$ were counted and the count was placed in bin number $n$; this count will be denoted $D_n$; the set of counts $D_n$ represents the experimental data. The theoretical model to be fit to the experimental data will be denoted $M_n$,

$$M_n=S(t_{n-1})-S(t_n) \quad (2)$$

Alternatively, one could define the experimental data $E_n$ as the number of dwell times greater than $t_n$ and then use $S(t_n)$ for the theoretical model. At first glance the two approaches appear to be equivalent, however, there is a significant difference in terms of error statistics. The random errors in the binned data $D_n$ are not correlated, which is a mandatory requirement for using the $\chi^2$ goodness-of-fit test and least-squares estimation, while the random errors in $E_n$ are highly correlated, which makes them unsuitable for statistical analysis.

In the case where all the bins are of equal width, the theoretical model can be expressed directly as a linear combination of exponential functions,

$$M_n = \sum_{i=1}^{N_{\exp}} \alpha_i \exp\left(-n\Delta t/\tau_i\right) \tag{3}$$

where

$$\alpha_i = \left[\exp\left(\Delta t/\tau_i\right) - 1\right] A_i \tag{4}$$

A weighted least squares estimator (LSE) was used to fit the model function based on equations (1-2) to the experimental data. LSE finds a set of parameters that minimizes the sum

$$SSQ = \sum_{n=1}^{N_D} W_n \left[D_n - M_n\left(\vec{p}\right)\right]^2 \tag{5}$$

where $N_D$ is the total number of data points, i. e., the number of bins in our case; $W_n$ is the weight for bin $n$, and $\vec{p}$ is the parameter vector, i. e., a set of all $A_i$ and $\tau_i$. The choice of the weights $W_n$ affects the accuracy of the parameter estimates $\vec{p}$ obtained by LSE. Hamilton has proven that the most accurate parameter estimates are obtained when the weights $W_n$ are equal to the inverse variances of the data $D_n$ [Walter Clark Hamilton *Statistics in physical science: Estimation hypothesis testing, and least squares* Ronald Press, New York, 1964],

$$W_n = \frac{1}{Var\left(D_n\right)} \tag{6}$$

With this weighting the *SSQ* defined in equation (5) becomes identical to the $\chi^2$ defined as

$$\chi^2 = \sum_{n=1}^{N_D} \frac{\left[D_n - M_n\left(\vec{p}\right)\right]^2}{Var\left(D_n\right)} \tag{7}$$

This makes it possible to apply the chi-squared goodness-of-fit test directly to the result of the *SSQ* minimization by the LSE. However, all this is contingent on our ability to accurately estimate the variances.

Since the counts $D_n$ represent the numbers of independent events, the numbers $D_n$ must obey Poissonian statistics. Thus, the variance of $D_n$ must equal the ensemble mean of $D_n$:

$$Var\left(D_n\right) = \left\langle D_n \right\rangle \tag{8}$$

The problem is that the ensemble means $\langle D_n \rangle$ are unknown. For the bins with very large counts $D_n$ the difference between $D_n$ and its ensemble mean is rather small, which makes it possible to substitute $D_n$ for $\langle D_n \rangle$ in those bins. For the bins with not very large counts $D_n$ such a substitution would result in a significant error, and for the bins with zero counts ($D_n$=0) this substitution would result in a division-by-zero error in equation (7). Clearly, substituting $D_n$ for $\langle D_n \rangle$ does not always work. Note, that if a model adequately fits the data, then the fit $M_n$ is closer to $\langle D_n \rangle$ than the data $D_n$, which contain random noise. The so-called exponential series model is ideally suited for this purpose because (i) it adequately fits any kinetic data and (ii) it is linear in all the adjustable model parameters, therefore the algorithm converges after just one iteration. The exponential series model is based on equations (1,2), where $N_{\exp}$ is very large; the time constants $\tau_i$ are equally spaced on the logarithmic scale and fixed, with $\tau_1$ being equal to $\Delta t$,

$\tau_{i+1}/\tau_i$=1.8, and $\tau_{N\text{exp}}$ being close to $N_D\Delta t$. The best fit $M_n$ achieved using the exponential series model is used for $Var(D_n)$ in equation (7). This approach has been extensively tested using simulated dwell time data and found to produce reliable $\chi^2$ values (suitable for applying the goodness-of-fit test) if only the data bins with $Var(D_n) \geq 0.1$ were included in the data analysis. If the data set included a tail where the exponential series fit resulted in $M_n$ significantly lower than 0.1, then the $\chi^2$ values were not always reliable and the goodness-of-fit test could lead to erroneous conclusions. For that reason, after applying the exponential series model to the full data set, the number $N_D$ was adjusted if necessary to exclude the tail with $Var(D_n)<0.1$.

The $\chi^2$ value calculated using equation (7) is not reduced; the reduced $\chi^2$ equals $\chi^2/(N_D-N_P)$, where $N_P$ is the number of free fitting parameters; if all $A_i$ and $\tau_i$ are free fitting parameters, then $N_P=2N_{\text{exp}}$. The Akaike Information Criterion (AIC) was calculated as AIC=$2N_P+\chi^2-N_D$ . Reduced $\chi^2$ and AIC values are shown in Suppl. Tables S1 and S2.

Binning of the dwell times, calculation of the weights, and fitting with the model function based on equations (1,2) was all carried out using the program Nexp_GLSE. The program is capable of global analysis, in which multiple data sets are analyzed simultaneously, with some of the parameters linked (shared) across multiple data sets, while other parameters are applied to just one data set. In addition to the minimized $\chi^2$ value and the best-fit estimates for the parameters the program also outputs the standard deviation for each parameter and the variance-covariance matrix for the random errors involved in the parameter estimates. The variance-covariance matrix can be used to estimate the standard deviation for any function φ of the parameters if the first partial derivatives of that function with respect to every parameter $p_i$ exist,

$$\sigma\varphi(\vec{p})=\sqrt{\sum_{i=1}^{N_P}\sum_{j=1}^{N_P} C_{i,j}\left(\frac{\partial\varphi}{\partial p_i}\right)\left(\frac{\partial\varphi}{\partial p_j}\right)} \tag{9}$$

where σφ is the standard deviation for the function φ and $C_{i,j}$ is an element of the variance-covariance matrix. This method was applied to calculate the standard deviations for the fractional amplitudes $f_i$ and the ensemble mean dwell time ⟨τ⟩, which were defined as follows:

$$f_i=\frac{A_i}{\sum_{j=1}^{N_{\text{exp}}} A_j} \tag{10}$$

$$\langle\tau\rangle=\frac{\sum_{i=1}^{N_{\text{exp}}} A_i\tau_i}{\sum_{j=1}^{N_{\text{exp}}} A_j} \tag{11}$$

Numerical values of the fractional amplitudes $f_i$ and the mean dwell time ⟨τ⟩ together with their standard deviations are presented in Figure 3 and in the SI tables S3B and S4.

**Suppression of systematic errors**

The program Nexp_GLSE was extensively tested using simulated dwell time data and was found to produce adequate fits (judged by the reduced $\chi^2$ values) in all tests without exception. However, when fitting binned experimental dwell times, the fit was not always adequate. Specifically, a statistically unacceptable deviation was often observed in bin number 1. Note, that bin number 1 contains those dwell times that do not exceed one video frame time (100 ms). For such a short dwell time the fluorescence intensity integrated over the frame time often falls below the detection threshold, therefore the number of dwell times in bin number 1 was underestimated, resulting in a systematic error. To minimize the effect of this systematic error the data for bin number 1 was excluded from the analysis for all data sets. This solved the problem with inadequate fits (apparently, this systematic error did not significantly affect the data in bins number 2 and above). Exclusion of bin number 1 was practically accomplished by setting its weight $W_1$ to zero instead of $1/Var(D_1)$; bins with zero weights were not included in the total count of data bins $N_D$ that was used in calculating the reduced $\chi^2$ and AIC.

**Worm-like chain simulation procedure**

Worm-like chain (WLC) simulations are employed in this work to produce a qualitative explanation of the dwell time dependence on the length of the nascent chain. A coarse-grained model represents the polypeptide as a chain consisting of segments $\vec{r}_i$, $i$=1,2,...,$N$, where $N$ is the number of amino acids (not counting those inside the ribosome exit tunnel and those that are part of the interaction motif defined on page 19 of this document). The length of each segment $l$ equals 0.36 nm and represents the contour length per one residue; the full contour length being equal to $Nl$. The persistence length $P$ equals 0.50 nm. Unconstrained WLC simulations are conducted in empty space; constrained simulations do not allow the WLC to enter the half-space $z \leq 0$ that is occupied by the ribosome. Below is the description of the algorithm that generates a WLC with the specified parameters $N$, $l$ and $P$ using a random number generator.

The direction of segment $i$ is parallel to a unit vector $\vec{d}_i$ that is defined as

$$\vec{d}_i = \frac{\vec{r}_i}{|\vec{r}_i|} = \frac{\vec{r}_i}{l} \tag{12}$$

Worm-like chain model assumes that the bending energy is directly proportional to the sum over all segments of the dot products of the unit vectors $\vec{d}_{i-1}$ and $\vec{d}_i$:

$$U = -\varepsilon \sum_{i=1}^{N} \left(\vec{d}_{i-1} \cdot \vec{d}_i\right) \tag{13}$$

The dot product of two unit vectors equals cosine of the angle between them,

$$\left(\vec{d}_{i-1} \cdot \vec{d}_i\right) = \cos\theta_i \tag{14}$$

therefore the bending energy can be also expressed in terms of the angles $\theta_i$:

$$U = -\varepsilon \sum_{i=1}^{N} \cos\theta_i \tag{15}$$

The vector $\vec{d}_{i-1}$ and the angle $\theta_i$ are insufficient to uniquely define the direction of the unit vector $\vec{d}_i$ : one more angle $\varphi_i$ is needed. One can think of $\theta_i$ and $\varphi_i$ as the bond angle and the dihedral angle (although the bond angle actually equals $\pi$-$\theta_i$). According to Boltzmann distribution, at an absolute temperature $T$ the probability density of the angles $\theta_i$ and $\varphi_i$ is an exponential function of the bending energy,

$$p(\theta_i,\varphi_i)\,d\theta_i\,d\varphi_i = C\,\exp\left(\frac{\varepsilon\cos\theta_i}{k_B T}\right)\sin\theta_i\,d\theta_i\,d\varphi_i \tag{16}$$

where $C$ is a normalization factor, $k_B$ is the Boltzmann constant, and $\sin\theta_i\,d\theta_i\,d\varphi_i$ is an elementary solid angle. Since the expression on the right-hand side of equation (16) is independent of $\varphi_i$, both sides of this equation can be integrated from 0 to $2\pi$ by $d\varphi_i$, yielding the probability density of the angle $\theta_i$:

$$p(\theta_i)\,d\theta_i = 2\pi C\,\exp\left(\frac{\varepsilon\cos\theta_i}{k_B T}\right)\sin\theta_i\,d\theta_i \tag{17}$$

The normalization factor $C$ can be calculated by equating to 1 the integral from 0 to $\pi$ of the right-hand side of Equation (17); substituting the resulting value of $C$ in equation (17) results in

$$p(\theta_i) = \frac{\xi\exp(\xi\cos\theta_i)\sin\theta_i}{\exp(\xi)-\exp(-\xi)} \tag{18}$$

where $\xi$ is a combination of three parameters,

$$\xi=\frac{\varepsilon}{k_B T} \tag{19}$$

Ensemble mean $\langle\cos\theta_i\rangle$ can be obtained by integrating $\cos\theta_i\,p(\theta_i)$ from 0 to $\pi$, which gives

$$\langle\cos\theta_i\rangle=L(\xi) \tag{20}$$

where $L(\xi)$ is the well known Langevin function, $L(\xi)=\coth(\xi)-1/\xi$, and $\coth(\xi)$ is a hyperbolic cotangent. According to the worm-like chain model the ensemble mean $\langle\cos\theta_i\rangle$ must also satisfy the following relation:

$$\langle\cos\theta_i\rangle=\exp(-l/P) \tag{21}$$

Combining equations (20) and (21) we obtain

$$L(\xi)=\exp(-l/P) \tag{22}$$

Resolving equation (22) with respect to $\xi$ allows one to express the new parameter $\xi$ in terms of the known parameters $l$ and $P$. In practice this requires calculating the inverse Langevin function, which can be accomplished using Newton-Raphson method (this is also a challenging task because both the Langevin function and its first derivative experience catastrophic cancellation in the vicinity of $\xi$=0). A Fortran subroutine for calculating the inverse Langevin function with the full precision of REAL(8) numbers was used.

The actual algorithm for simulating a WLC consists of a few preparatory operations followed by *N* steps during which the WLC is gradually built up one amino acid at a time. The preparatory operations consist of: (i) calculating the parameter ξ from the parameters *l* and *P* as described above, (ii) defining the initial direction $\vec{d}_0=(0,0,1)$ which is normal to the surface of the ribosome and pointing outside of the exit tunnel, and (iii) defining the radius-vector $\vec{R}_0=(0,0,0)$ that defines the coordinates of the N-terminal of a zero-length nascent chain.

Algorithm step number *i* (*i*=1,2,...,*N*) consists of the following operations. First, two unit vectors are constructed that are both normal to $\vec{d}_{i-1}$ and also normal to each other. Toward this goal the algorithm finds the component of the vector $\vec{d}_{i-1}$ that is smallest in absolute value and constructs a unit vector $\vec{e}_{i-1}$ parallel to this component. For example, if the *x* component of vector $\vec{d}_{i-1}$ is the smallest in magnitude, then $\vec{e}_{i-1}=(1,0,0)$ , if the *y* component of vector $\vec{d}_{i-1}$ is the smallest in magnitude, then $\vec{e}_{i-1}=(0,1,0)$ , etc. After that the vector $\vec{e}_{i-1}$ is orthogonalized with the unit vector $\vec{d}_{i-1}$ and normalized to a unit length:

$$\vec{f}_{i-1}=\frac{\vec{e}_{i-1}-(\vec{d}_{i-1}\cdot\vec{e}_{i-1})\vec{d}_{i-1}}{\left|\vec{e}_{i-1}-(\vec{d}_{i-1}\cdot\vec{e}_{i-1})\vec{d}_{i-1}\right|} \tag{23}$$

Vector $\vec{f}_{i-1}$ is a unit vector and it is normal to $\vec{d}_{i-1}$ . The third unit vector $\vec{g}_{i-1}$ that is normal to both $\vec{d}_{i-1}$ and $\vec{f}_{i-1}$ is calculated as the cross product

$$\vec{g}_{i-1}=\left[\vec{d}_{i-1}\times\vec{f}_{i-1}\right] \tag{24}$$

Next a random value for cos$\theta_i$ is generated

$$\cos(\theta_i) = \frac{1}{\xi}\,\ln\left\{\,\exp(\xi) - \left[\exp(\xi)-\exp(-\xi)\right]*\mathrm{rand}()\,\right\} \tag{25}$$

where rand() is a pseudo-random number generator in Fortran programming language, this function produces pseudo-random values uniformly distributed between 0 and 1. It can be shown that the cos$\theta_i$ values generated using equation (25) strictly comply with the probability distribution in equation (18). The value of sin$\theta_i$ is calculated as

$$\sin(\theta_i)=\sqrt{1-\cos^2(\theta_i)} \tag{26}$$

Next a random value for $\varphi_i$ is generated; this angle is uniformly distributed between -π and +π.

$$\varphi_i=2\pi\left[\mathrm{rand}()-\frac{1}{2}\right] \tag{27}$$

Next the new unit vector $\vec{d}_i$ is calculated, followed by the new vectors $\vec{r}_i$ and $\vec{R}_i$ :

$$\vec{d}_i=\cos(\theta_i)\vec{d}_{i-1}+\sin(\theta_i)\cos(\varphi_i)\vec{f}_{i-1}+\sin(\theta_i)\sin(\varphi_i)\vec{g}_{i-1} \tag{28}$$

$$\vec{r}_i=l\vec{d}_i \tag{29}$$

$$\vec{R}_i=\vec{R}_{i-1}+\vec{r}_i \tag{30}$$

In the case of unconstrained simulations this represents the last operation of step number *i*. Constrained simulations include one more operation that will be discussed later.

The worm-like chain model (unconstrained) predicts the following correlation between the direction $\vec{d}_i$ and $\vec{d}_{i+m}$ :

$$\langle(\vec{d}_i \cdot \vec{d}_{i+m})\rangle = \exp(-ml/P) \quad (31)$$

This relation must hold for any values of *i* and *m*. To test the algorithm of unconstrained WLC simulations described above, $10^8$ WLCs were simulated, and the mean values of the dot products of vectors $\vec{d}_i$ and $\vec{d}_{i+m}$ were calculated for a wide range of *m* values. It was confirmed that the relation in equation (31) holds with great accuracy (the accuracy is determined by the number of WLCs simulated). This convinced the authors that the algorithm works exactly as expected.

In the case of constrained WLC simulations equation (31) is not expected to hold, however, all the operations that constitute the algorithm step number *i* are still applicable and justified. The radius-vector $\vec{R}_i$ generated at the end of step *i* consists of three components: $X_i$, $Y_i$, $Z_i$. The component $Z_i$ must be positive for all $i$=1,2,...,$N$. If at the end of the step number *i* the value $Z_i$ is negative or equals zero, then this WLC is not allowed. Two actions can be considered in this case. If the nascent chain is considered to be rigid so that it would rather move as a whole than change its conformation, then upon encountering $Z_i \leq 0$ the algorithm should be restarted at $i$=1 and a new WLC should be generated from the start. However, if the nascent chain is considered to be flexible so that it would rather change the angles $\theta_i$ and $\varphi_i$ than move away from the surface of the ribosome, then upon encountering $Z_i \leq 0$ the algorithm should be sent back just one step to $i$=$i$-1 and repeat the last step as many times as it takes until $Z_i$>0 is obtained. The latter approach was used in all simulations reported in this work.

The number of amino acids denoted as *L* in Figure 5 A, B or as "NC length" in Figure 5 C differs from *N* for two reasons: the first 32 amino acids counting from the peptidyl-transferase center (PTC) are located inside the ribosome exit tunnel, whereas the last 10 amino acids comprise the interaction motif (see Suppl. Fig. S6 panel C). Thus, $L$=32+$N$+10=$N$+42; $N$=$L$-42. Red spheres in Figure 5 A denote the position of amino acid number 11 counting from the N-terminus. The distance *R* in Figure 5 B is measured from the ribosome tunnel exit to amino acid number 11 counting from the N-terminus. In Figure 5 C $c_{\text{eff}}$ is the concentration of amino acid number 11 counting from the N-terminus.

To generate the data for Figure 5 B, $10^8$ WLCs were simulated for each of the six *N* values; the lengths of the vectors $\vec{R}_N$ were binned using bin widths $\Delta R$=0.04 nm. The probability density was calculated as the number of counts in a bin divided by the product ($\Delta R$ times the total number of WLCs).

To generate the data for Figure 5 C, we utilized the symmetry of the problem due to which the local concentration of WLC ends depends on only two variables: $P=\sqrt{X^2+Y^2}$ and *Z*. The values of *P* and *Z* were calculated for each of the four TF binding sites denoted a, b, c, d, thus yielding $P_a$, $Z_a$, $P_b$, $Z_b$, $P_c$, $Z_c$, $P_d$, $Z_d$. $10^8$ WLCs were simulated, and at each simulation step *i* the vector $\vec{R}_i$ was used to calculate $P_i$ and $Z_i$. $N$=258 bins were created for each of the sites a, b, c, d. If the condition $(P_i-P_a)^2+(Z_i-Z_a)^2 \leq (\Delta r)^2$ was satisfied, then a count was added to bin number $i$ for site a. Likewise, if the condition $(P_i-P_b)^2+(Z_i-Z_b)^2 \leq (\Delta r)^2$ was satisfied, then a count was added to bin number $i$ for site b. The cutoff distance $\Delta r$ was chosen equal to 0.2 nm. The local concentration was calculated as the number of counts in a bin divided by the product (toroid volume times the total number of WLCs), where the toroid volume for site a was calculated as $2\pi^2 P_a(\Delta r)^2$, for site b as $2\pi^2 P_b(\Delta r)^2$, etc.. To convert concentration from molecules per $nm^3$ to moles per liter, it was multiplied by $10^{24}$ $nm^3$/liter and divided by the Avogadro number.
